# The Hybrid Drive: a chronic implant device combining tetrode arrays with silicon probes for layer-resolved ensemble electrophysiology in freely moving mice

**DOI:** 10.1101/2021.08.20.457090

**Authors:** Matteo Guardamagna, Ronny Eichler, Rafael Pedrosa, Arno Aarts, Arne F. Meyer, Francesco P. Battaglia

**Author notes:** Correspondence (M.G.), (F.P.B.). Co-senior author (Arne F. Meyer, Francesco P. Battaglia).

## Abstract

Understanding the function of brain cortices requires simultaneous investigation at multiple spatial and temporal scales and to link neural activity to an animal’s behavior. A major challenge is to measure within- and across-layer information in actively behaving animals, in particular in mice that have become a major species in neuroscience due to an extensive genetic toolkit. Here we describe the Hybrid Drive, a new chronic implant for mice that combines tetrode arrays to record within-layer information with silicon probes to simultaneously measure across-layer information. The design of our device combines up to 14 tetrodes and 2 silicon probes, that can be arranged in custom arrays to generate unique areas-specific (and multi-area) layouts. We show that large numbers of neurons and layer-resolved local field potentials can be recorded from the same brain region across weeks without loss in electrophysiological signal quality. The drive’s lightweight structure (*≈* 3.5 g) leaves animal behavior largely unchanged during a variety of experimental paradigms. We demonstrate how the data collected with the Hybrid Drive allow state-of-the-art analysis in a series of experiments linking the spiking activity of CA1 pyramidal layer neurons to the oscillatory activity across hippocampal layers. Our new device fits a gap in the existing technology and increases the range and precision of questions that can be addressed about neural computations in freely behaving mice.

## Introduction

A major goal in neuroscience is to understand the function of the nervous system at a variety of levels, from single cells to entire networks that mediate complex behaviors such as navigation, foraging, and social interaction in natural settings. The mouse has emerged as an important species in neuroscience due to the availability of genetic tools that can reveal the precise mechanisms at the circuit level and link these to the animal’s behavior (Luo et al., 2008). In particular, many brain structures (including the cortex and the hippocampus) are organized in layers with information processing within and also across the layers (Andersen et al., 1971, Amaral and Witter, 1989, Andersen et al., 2006). Simultaneously measuring neural activity at these two dimensions while an animal is interacting with its environment remains a major challenge, especially in the mouse with its relatively small size (body weight of an adult male mouse *≈* 25/30 grams). To address this challenge, we developed a chronic implant to measure neural activity within and across layers in freely moving mice.

Tetrodes are a standard method for neuronal recordings in behaving animals, in particular for chronic recordings of many neurons for extended time periods in freely moving rodents (McNaughton et al., 1983, Wilson and McNaughton, 1993, Gothard et al., 1996). The precision, stability and bio-compatibility of tetrode implants enable recordings for months from the same local circuit (Voigts et al., 2013, 2020). Modern tetrode implants enable highly flexible arrangements of tetrodes that can easily be modified depending on the set of target areas. By performing small and repeated adjustments of individual tetrodes, it is possible to maximize cell yield and to target specific layers (Harris et al., 2000) or structures deep in the brain (Chen et al., 2020) without the need to remove overlying structures as for optical imaging methods. However, a disadvantage of tetrodes is that they do not easily allow for measuring the activity across layers with a well-defined geometrical arrangement of recording sites. High-density silicon probes (Buzsáki et al., 2015), on the other end, offer unparalleled opportunities for populations recordings across several layers or even from multiple brain regions in head-fixed (Steinmetz et al., 2019) or freely moving animals (Steinmetz et al., 2021). Their geometric design – electrodes are typically arranged along one or more shafts – enables superior anatomical localization, given the well-defined distances between recording sites. Yet, silicon probes lack the flexibility of tetrodes arrays which can easily be arranged in unique geometric layouts to sample different locations along a single layer (or adjacent areas). While custom silicon probe designs are possible, they are typically expensive and tailored to a single brain structure.

Ideally, one would combine in a single device the flexibility and sampling capability of custom tetrode arrays, to record neural activity within small layers, and silicon probes to record activity across layers and structures. Microdrives combining tetrodes and silicon probes have been developed for rats (Headley et al., 2015, Michon et al., 2016). However, mouse-specific designs are strictly limited by the weight and the available space on the head of the animals. Here we describe the Hybrid Drive, a novel chronic implant that combines the advantages of tetrodes and flexible-shaft silicon probes in a single, lightweight device. A core feature of the Hybrid Drive is its ability to pack up to 14 tetrodes and 2 silicon probes in small arrays. The design of the array can easily be adjusted depending on the anatomical or functional organization of the target area (e.g., the proximo-distal organization of CA1 or the tonotopic organization of sensory cortices). The array can target a single area, two adjacent areas or even be divided into two or more bundles to target distant regions. Similarly, the choice of the contact displacement of the flexible-shaft silicon probe can be adapted to match the three-dimensional organization of the desired brain areas. Importantly, the overall weight of the implant (*≈* 3.5 grams) enables studies in freely moving animals, including mice.

We show that the ability of the Hybrid Drive to record single cells with tetrodes is not affected by the presence of a silicon probe (and vice versa) and patterns of behavior are largely unchanged with respect to other implant device. We further demonstrate the potential of the Hybrid Drive in a series of experiments in freely moving mice, focusing on recordings from the hippocampal CA1 region. First, we show that the interactions of single cells and layer-specific local field potentials in CA1 strongly vary with the animal’s behavior, across slow and fast time scales. Second, we show that single cells in the hippocampal pyramidal layer integrate across-layer information, in addition to spatial variables typically studied in navigation experiments. This integration is layer-specific, is consistent across different animals, and improves the prediction accuracy of the cells’ responses. These results already demonstrate that the Hybrid Drive can lead to novel insights into the interactions between neural processes occurring within and across layers and the animal’s behavior.

## Results

### A chronic implant for simultaneous measurements of within-layer and across-layer neural activity in freely moving mice

The Hybrid Drive consists of a custom chronically implantable tetrode drive (based on an existing implant design; Voigts et al. (2013)) which can include up to two silicon probes (Figure 1A). The 14 independently movable tetrodes enable flexible recordings of local activity, e.g., from neurons within the same cortical or hippocampal layer, while the two movable silicon probes provide across-layer information. In order to tightly pack the tetrodes and the silicon probes in a single drive body and polymide tube array we developed a new custom electrode interface board (EIB; Figure 1D) combining two lateral ZIF connectors (16 channels each) and 56 pin holes for wire electrodes - integrated with three 36-pins Omnetics connectors.

**Figure 1:**
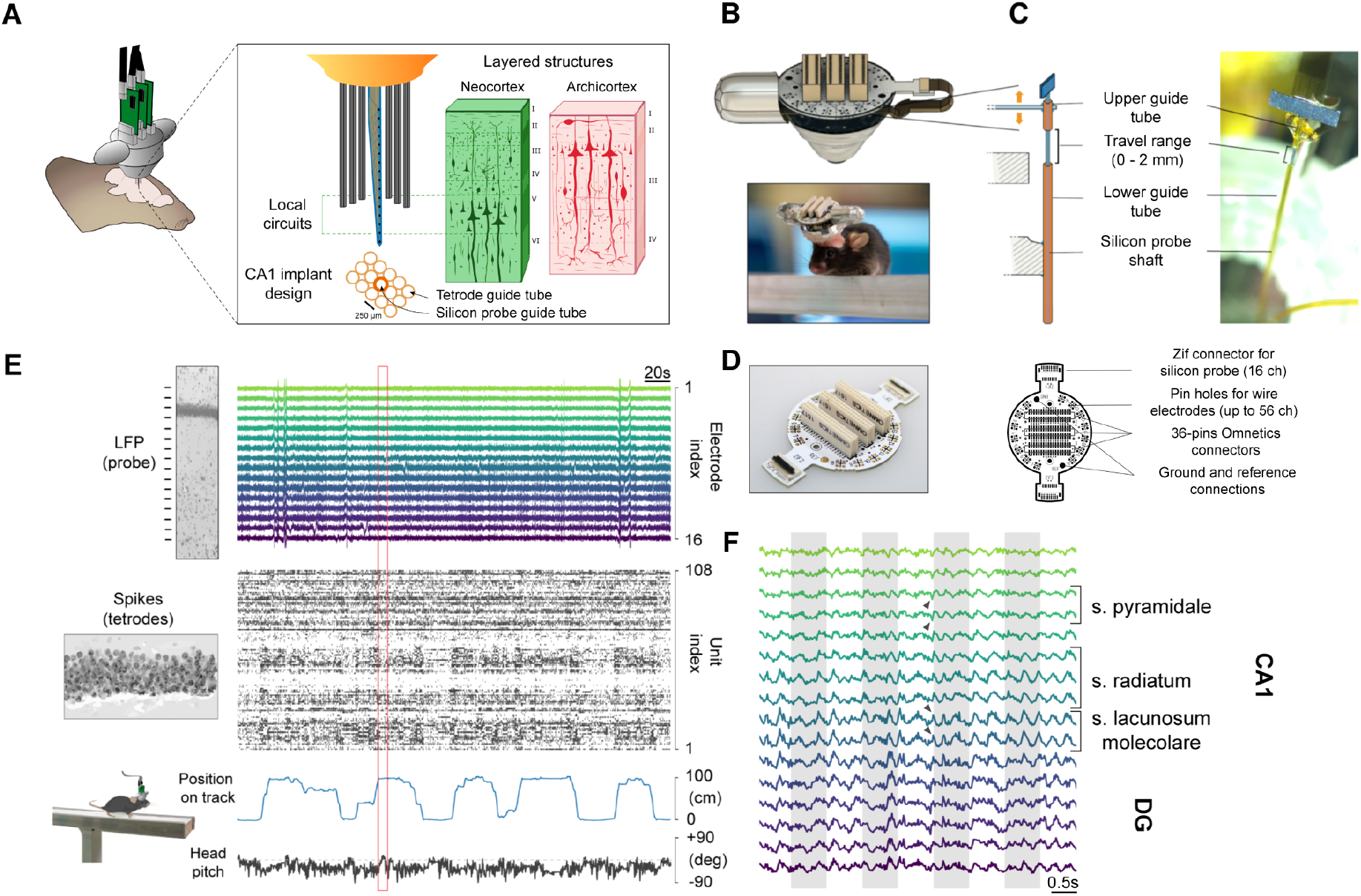
Simultaneous measurement of within-layer and across-layer electrophysiological activity in a freely moving mouse. (A) Neural activity is recorded using a chronic implantable “Hybrid Drive” that contains 14 tetrodes and 1–2 silicon probes. The silicon probes record neural activity across layers of a given structure, while tetrodes can record activity of single cells from within a single layer. Example 5 x 3 array design for dorsal CA1 recordings. Mouse brain illustration adapted from SciDraw (doi.org/10.5281/zenodo.3925919). (B) 3D model of a fully-assembled Hybrid Drive. Lateral plastic cone shields protect the flex cables and ZIF connectors of the silicon probes. (C) Left, illustration of the spring-screw mechanism for independently movable silicon probes and tetrodes. Right, example photo of a silicon probe positioned in the Hybrid Drive. Bottom, photo of the Hybrid Drive implanted in a 5 month old C57BL6/J mouse. (D) Left, EIB picture. Right, illustration of the EIB design. The two lateral ZIF connectors for the silicon probes are integrated with the 36-pins Omnetics connectors in a compact design. 56 pin holes for wire electrodes allow to include up to 14 tetrodes. Ground and reference connections are positioned in between the ZIF and Omnetics connectors. (E) Example traces of neural and behavioral variables simultaneously recorded with the Hybrid Drive in an animal traversing a linear track. Top: Local field potential (LFP) recordings from the silicon probe, spanning dorsal CA1. Middle: raster plot of single units recorded with tetrodes in the CA1 pyramidal layer. Bottom: head pitch and the animal’s position on the linear track. (F) Zoomed-in LFP traces for the region marked by the red rectangle in E. LFP traces show typical markers of CA1 layers, including depth-dependent oscillation amplitude and phase reversal in deeper layers (highlighted by dark grey arrows). Shaded gray regions for easier visualization of temporal alignment.

The use of Omnetics connectors on the EIB ensures compatibility with most modern neural recording systems, including the Open Ephys acquisition system (Siegle et al., 2017). Another unique feature of the implant is the flexible design of the guide array holding the tetrodes and the silicon probes (Figure 1A). When choosing the target area of interest the user should carefully plan the geometrical displacement of the tetrode tube arrays. Briefly, each tube segment should be stacked and glued together to obtain the desired shape (Figure 1A); 5×3 rectangle). Subsequently, the whole guide tube array can be glued at a specific angle to the drive body. This is possible thanks to the silicon probe shaft flexibility, a key aspect of the Hybrid Drive’s design (Herwik et al., 2011). It facilitates the insertion of the probe in the guide array during drive building (Figure S1) and provides additional protection from mechanical shocks, e.g., in the animal’s home cage or during experiments. Thanks to these features the position of the silicon probes relative to the tetrodes, or the angle of the overall assembly, can easily be modified depending on brain areas under investigation (Figure 2). The position of the tetrodes and the silicon probes in the guide tube array can be mapped during drive building and, at the end of the experiments, it can be matched to the final recording location in Nissl-stained brain sections (Figure 2B). In the current design, we used one or two 16-channel probes (E16R_60_S1_L20_NT with 60 µm pitch between contacts, ATLAS Neuroengineering, Belgium), optimized for long-term recordings in the hippocampus, a brain region that is widely being studied in freely moving rodents. The weight of the whole assembly is *≈* 3.5 g. This enables the study of naturalistic behaviors, such as navigation, exploration, and social interactions in unrestrained animals. Mice implanted with a Hybrid Drive targeting the hippocampal CA1 region readily explored the environment while enabling recordings from large numbers cells in the CA1 pyramidal layer together with local field potentials (LFPs) from all CA1 layers (Figures 1E,F) during a variety of behavioral paradigms.

**Figure 2:**
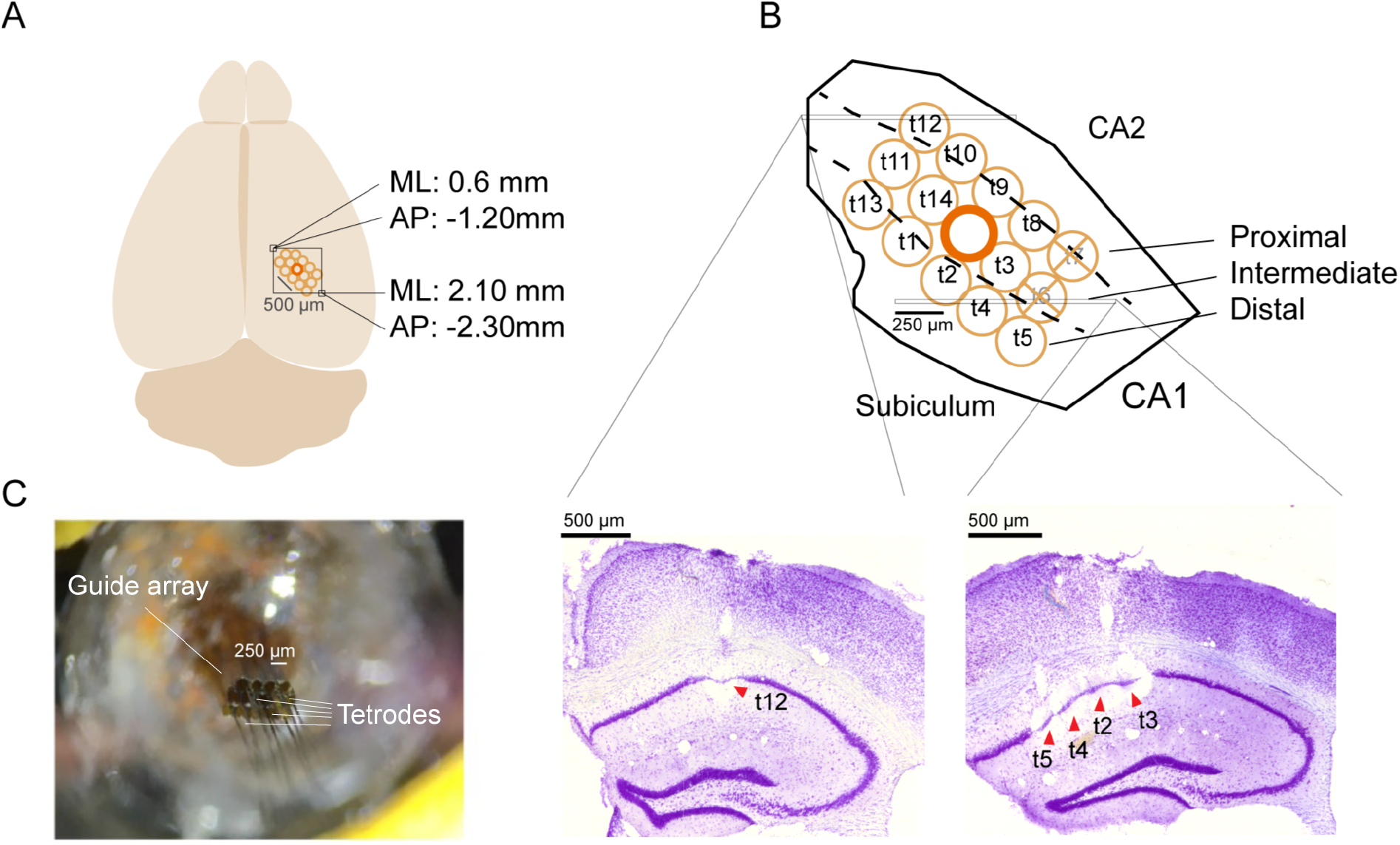
The array design can be adapted to match the shape and the organization of the target brain area. (A) Dorsal CA1 design. 5×3 tube array covering the dorsal CA1 area along its proximo-distal axis. Coordinates for the elliptic craniotomy are: top-left corner at AP: −1.20 mm; ML: 0.6 mm relative to bregma; bottom-right corner at AP: −2.30 mm; ML: 2.10 mm relative to bregma. The guide tube array is represented in orange, with the mapping of individual tetrodes (and silicon probe) over the target structures. The silicon probe (represented by the thicker tube in the image) is designed to cross all CA1 layers, while the tetrodes target the pyramidal layer. Brain illustration adapted from SciDraw (doi.org/10.5281/zenodo.3925941). (B) Organization of the bundle array over the CA1 region of the hippocampus. The bundle has been design to cover and sample as much as possible along CA1 proximo-distal axis. The position of individual tetrodes (and silicon probe) is mapped within the guide array during drive building. Then, individual tetrode positions can be traced back in the target area after histological procedures. Bottom, Nissl-stained coronal section showing recording locations in a more anterior brain slice (where the electrolytic lesion of t12 is visible) and a more posterior one (where the electrolytic lesions of t5,t4,t3,t2 are visible). t6 and t7 never reached the CA1 pyramidal layer. (C) Picture of the bottom array before cutting and plating. All tetrodes are drove out of the guide array, while the silicon probe shaft is retracted to avoid damage during the cutting procedure.

### The Hybrid Drive enables stable recordings of neural activity across weeks

We first checked that the presence of the silicon probe did not interfere with the quality of concomitant tetrode recordings (and vice versa; Figure 3). To quantify recording quality, the tetrodes of the Hybrid Drive were implanted in the pyramidal layer of dorsal CA1 while the linear silicon probe spanned the entire CA1 region, orthogonal the pyramidal layer (Figure 1E). Recordings were performed in three animals in a variety of tasks across at least 20 days. For each animal, experimental day 1 started when the majority of tetrodes exhibited sharp wave ripple (SWR) complexes during rest sessions in the home cage or clearly discernible theta oscillations during locomotion.

**Figure 3:**
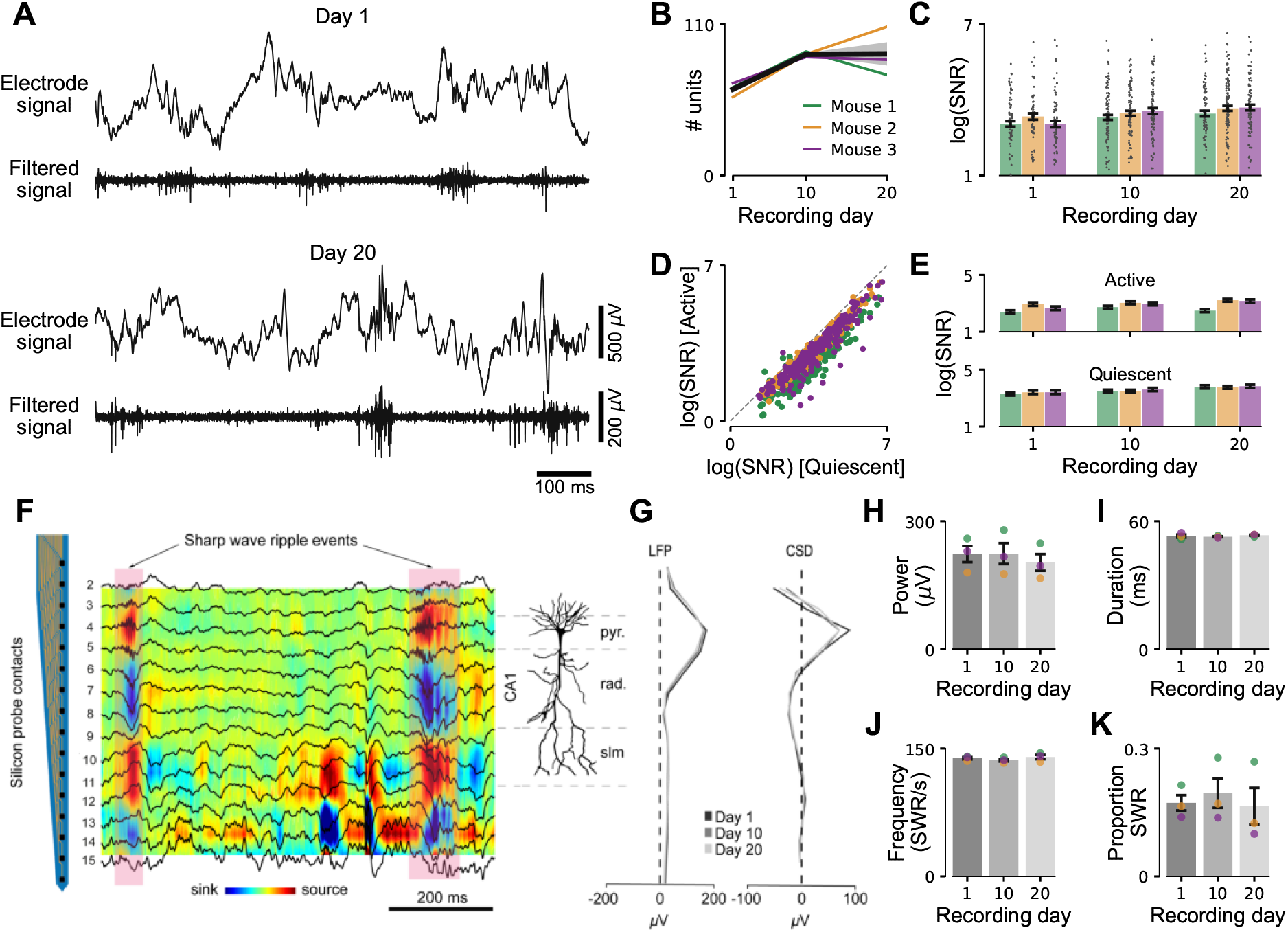
Neural recording quality is not affected when combining tetrodes with silicon probe recordings. (A) Broadband electrode signal from an example tetrode channel and high-pass filtered signal for the same channel on the first recording day (day 1; top) and 19 days later (day 20, bottom). (B) Number of isolated single units from 13 tetrodes positioned in the CA1 pyramidal layer for different recording days (mean day 1: 62.6 ± 5.1, mean day 10: 88 ± 2, mean day 20: 88.3 ± 17.8). Colors indicate different animals. (C) Signal-to-noise ratio (SNR) for different recording days. Gray dots indicate SNR estimates for single units. Same data as in B. Mean ± SEM. (D) SNR for different behavioral states (Active, Quiescent) pooled across days. Same data as in B. (E) The same as in C but separated into Active and Quiescent states. Increases in SNR in the presence of silicon probes can not be explained by changes in behavior across days. (F) Current-source density (CSD) computed from silicon probe targeting different CA1 layers during a rest session in the animal’s homecage. Superimposed LFP signals are aligned with CSD sink and sources. Silicon probe channels shown to the left. Example sharp wave ripple (SWR) events indicated by shaded red areas. (G) High-frequency oscillation (100-250 Hz) power profiles along the silicon probe for LFP (left) and CSD (right) signals. The profiles remain stable across days (Day 1, 10 and 20) with a clear peak in the pyramidal layer electrode. (H-K) SWR power, SWR duration, SWR frequency, and proportion of SWR events across recording days (during quiescent state). One home cage rest session per animal (each approx. 60 min).

Single units could consistently be recorded across weeks (Figure 3B). The number of isolated units was slightly lower on the first recording days, potentially resulting from uncertainty in positioning the tetrode tips in the pyramidal layer. Fine adjustments of the depths of individual tetrodes enabled the precise placement of tetrodes in the pyramidal layer (*≈* 80 microns thickness) to maximize the yield of simultaneously recorded cells.

To test the quality of detected spikes, we estimated signal-to-noise (SNR) measures (Meyer et al., 2018, Magland et al., 2020) for each tetrode channel for different recording days. Across all three animals, SNR increased across days (Figure 3C). We wondered if these changes in SNR across days can be explained by changes in behavior. To test this, we segmented the data into “Active” and “Quiescent” periods based on signals measured using the head-mounted accelerometer (Meyer et al., 2018) and computed SNR values for each condition. While cells had a lower SNR during the “Quiescent” compared to the “Active” condition (Figure 3D), this difference in SNR could not explain changes in SNR within conditions across days (Figure 3E). This suggests that the Hybrid Drive can reliably record neural ensembles for an extended period, similar to what is know for standard tetrode implants (Voigts et al., 2013, 2020).

Finally, we investigated common electrophysiological markers to assess the recording quality of the chronically implanted silicon probe. We focused on sharp wave ripple (SWR), distinct oscillatory patterns in the LFP often associated with memory-related processes during sleep (Buzsáki, 2015). Figure 3F shows unfiltered LFP signals exhibiting SWR events, together with the Current Sources Density (CSD) estimated from the LFP data. The SWR power depth profile maintained a consistent shape throughout the different weeks (Figure 3G, left). Similar results were found after applying CSD analysis to extract layer-independent oscillatory patterns (Figure 3G, right). The stability of the probe and the quality of the recorded signals were confirmed by multiple SWR measures. We found no significant differences in either SWR power (Figure 3H; One-way ANOVA, *p* = 0.82), SWR duration (Figure 3I; One-way ANOVA, *p* = 0.69), SWR frequency (Figure 3J; One-way ANOVA, *p* = 0.55) and proportion of detected SWR events (Figure 3J; One-way ANOVA, *p* = 0.86) between recording days 1, 10, and 20.

### Patterns of behavior are minimally disturbed by the drive implant

Next, we investigated the impact of the Hybrid Drive implant on gross locomotor and exploratory animal behavior. Previous work has shown that mice tolerate tetrode implants such as the flexDrive with only minimal changes in natural behavior (Voigts et al., 2013, Meyer et al., 2018, Voigts et al., 2020). To test if this is also true for the Hybrid Drive with the addition of silicon probes, we first analyzed locomotion patterns of mice traversing a linear track (Figure 4A). Animals received a reward at the ends of the track every time they completed a full lap (end-to-end run). Even without food deprivation, animals learned to consistently move from one end to the other to receive a reward; their running behavior became more stereotypical with increasing numbers of laps across days (Figure 4B). This increase was consistent across animals (Figure 4C) and was statistically significant in two of the three animals (Mouse 1, +2.1 laps per day, *p* = 0.03; Mouse 2, +2.6 laps per day, *p* = 0.004; Mouse 3, +1.7 laps per day, *p* = 0.48; Wald Test on slope of the linear fit). Mice were also running faster as indicated by the change in the running speed profile (Figure 4D; permutation test days 1–5 vs days 6–10, *p* = 0.029). We examined more closely the potential effects of the size and weight of the drive on the behavior of an example mouse, paying attention to head movements across days (Figure 4E). Starting from recording day 5 on the linear track, the mouse was holding its head closer to its natural position (typical pitch *≈* −25 deg; Meyer et al. (2018)). On day 10, we observed a decrease in variability, potentially resulting from more stereotypical running (day 1, pitch *−*34.9 *±* 14.4 deg, roll 2.2 *±* 8.7 deg; day 5, pitch *−*28.6 *±* 15.3 deg, roll 3.1 *±* 9.0 deg; day 10, pitch *−*28.7 *±* 11.5 deg, roll 2.7 *±* 9.5 deg).

**Figure 4:**
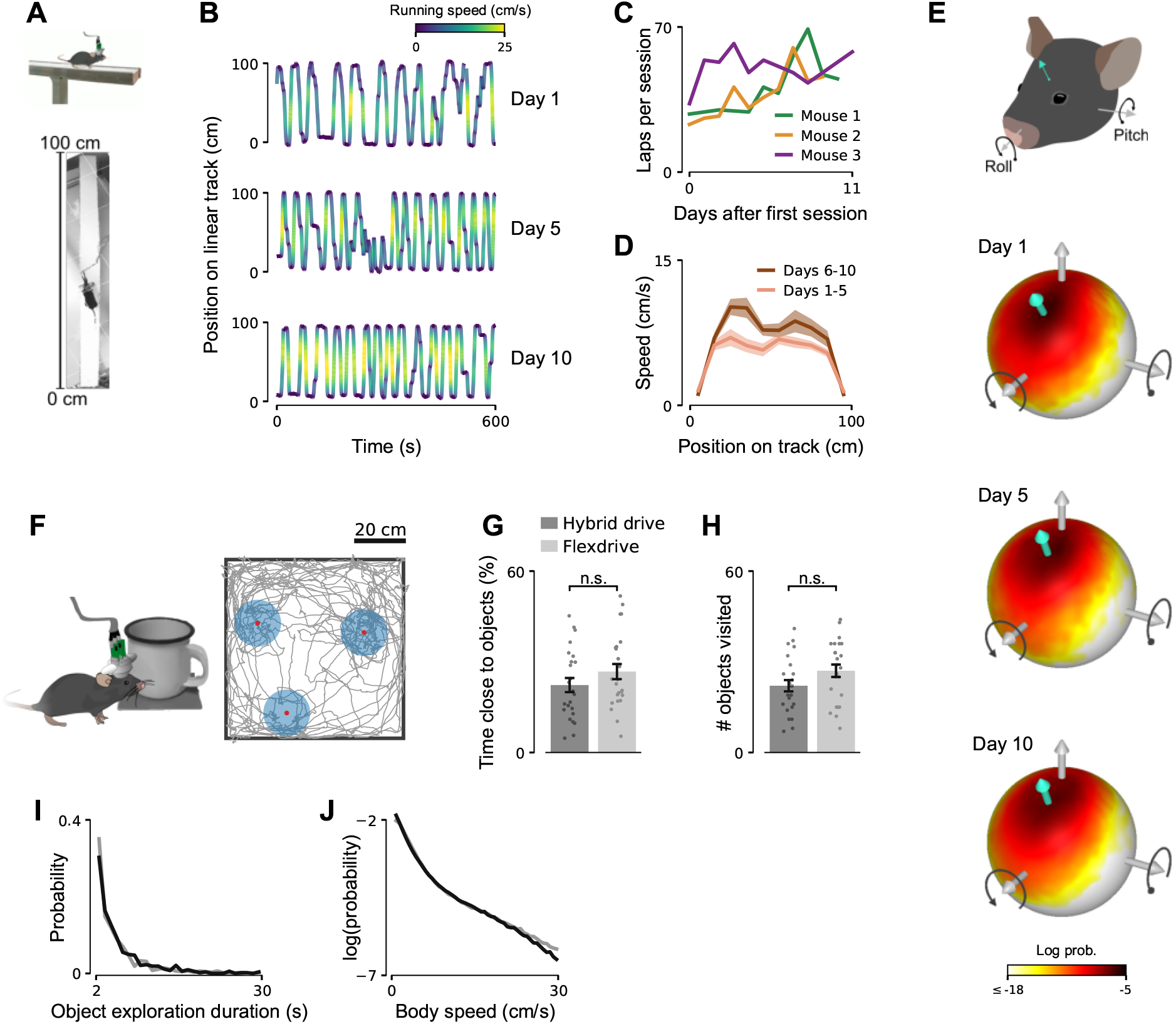
Patterns of behavior are minimally disturbed by the drive implant. (A) Mice implanted with a Hybrid Drive learned to traverse a linear track to receive rewards provided at the ends of the track. (B) 10-min example segments from different recording days (1, 5, and 10). Even without food deprivation, mice show increasingly stereotypical running behavior. Color denotes running speed. (C) Number of laps (end-to-end runs) per session. Data from three animals (n=9, 10, 10 sessions in mouse 1 (19.25 ± 4.04 minutes), 2 (29.15 ± 5.75 minutes), and 3 (25.17 ± 4.68 minutes), respectively). (D) Average running speed profiles for early (days 1–5, black line) and late (days 6–10, gray line) recordings. Mean ± SEM. Same data as in B. (E) Log-probability distribution of head tilt for day 1 (top), 5 (middle), and day 10 (bottom) for the data shown in C (mouse 1). Illustration indicates pitch and roll axes. Turquoise arrow indicates mean orientation of vertical (ventral-dorsal) head axis. (F) Object exploration task in which mice are freely exploring three objects in a square arena (width = 75cm). Red dots indicate object centers. Shaded blue circles (radius = 10 cm) show the area around each object that was considered as close to the object. Gray line indicates position of the mouse’s head in the arena for a 10-min trial. (G) Time close to objects for animals implanted with a Hybrid Drive (dark gray) or a flexDrive (light gray). Dots show single trials (10 min each). Number of trials: Hybrid Drive, n=7, 12, 5 trials in mouse 1, 2, 3, respectively; Flex drive, n=12, 6, 6 trials in mouse 4, 5, 6, respectively. Mean ± SEM. (H) Number of objects visited per trial. Mean ± SEM. Same data as in G. (I) Distribution of object exploration duration. Same data as in G. (J) Log-scaled distribution of body speed for animals implanted with a Hybrid Drive or flexDrive. Same data as in G.

Finally, we compared exploratory behaviors in two-dimensional environments that contained a small number of objects. Animals were free to explore a square environment (width: 75 cm) for 10 minutes. This was repeated on the same recording day to yield 5–7 consecutive trials per animal per day. We tracked the animal’s positions from camera recordings using a convolutional deep neural network trained using transfer learning (Mathis et al., 2018). To obtain baseline measures of animals with a slightly more lightweight neural implant (*≈* 2 grams) without silicon probes, we implanted three animals with a flexDrive (Voigts et al., 2013).

We found no difference in exploration patterns between animals implanted with a Hybrid Drive or a more lightweight flexDrive. Animals spent similar amounts of time close to objects (Figure 4G; Wilcoxon rank-sum test, *p* = 0.17) and visited a comparable number of objects during a trial (Figure 4H; Wilcoxon rank-sum test, *p* = 0.099). There was also no discernible difference in object exploration duration (Figure 4I; permutation tests, *p* = 0.17) and in body speed (Figure 4J; permutation tests, *p* = 0.23).

We conclude that learning of goal-directed behaviors and the ability to explore, and interact with, more complex environments were minimally affected by the presence and additional weight of our design.

### Interactions of spiking and across-layer neural activity vary with behavioral state in freely moving mice

We next explored the capacity of the Hybrid Drive to identify interactions between the spiking activity of single cells and across-layer LFP signals during different behaviors. Animals that can freely interact with their environments show a wide range of behaviors, including active exploration, quiescence, and sleep (Krakauer et al., 2017). To identify neural correlates of these behavioral “states”, mice implanted with a Hybrid Drive in CA1 were free to explore a rectangular open field arena (Figure 5A). Behavioral variables were measured using an external video camera and the head-mounted accelerometer. Before and after the open field recording, mice were allowed to naturally transition to sleep in their home cages while we recorded from the same cells as in the open field condition. Segmenting the open field recording into “Active” (body speed *≥* 2 cm/s) and “Quiescent” (body speed *<* 2cm/s) periods using video data allowed us to compare neural activity across four different conditions: “Pre sleep”, “Active”, “Quiescent”, and “Post sleep”. To exclude movement periods during the home cage sleep condition, we focused on periods during which mice were not moving their heads or bodies.

**Figure 5:**
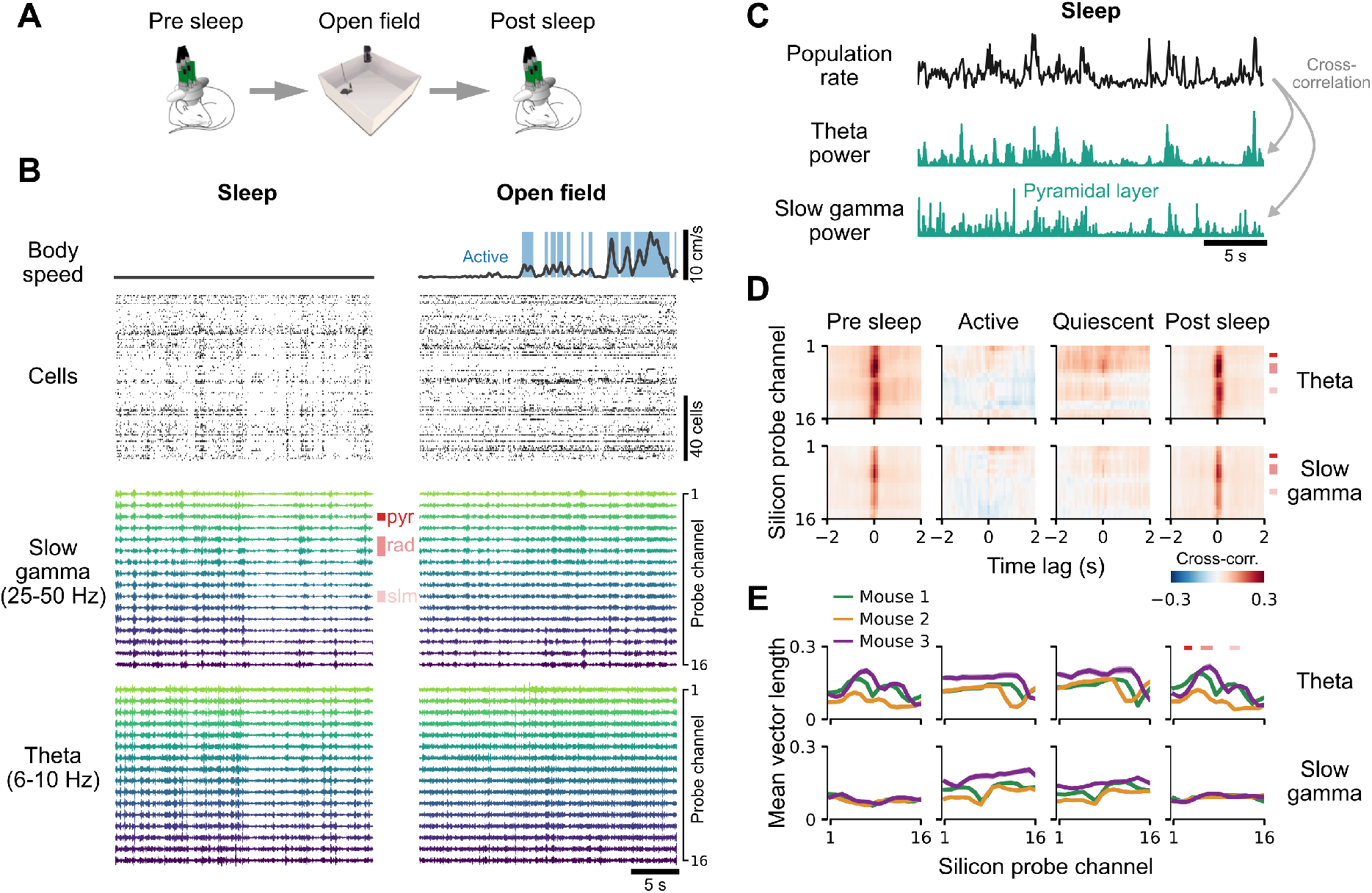
Interactions of spiking and across-layer neural activity vary with behavioral state in freely moving mice. (A) Neural activity and behavioral variables were measured while an animal was either sleeping or freely exploring an open field environment. (B) Example traces of simultaneously measured behavioral and neural variables for sleep (left) and open field exploration (right). Blue shaded regions in body speed trace indicate “Active” periods (body speed ≥ 2 cm/s). All other periods were considered as “Quiescent”. Vertical red bars indicate locations of pyramidal (pyr), radiatum (rad), and slm layers relative to the silicon probe electrodes. (C) Population rate and LFP power in the theta and slow gamma frequency bands for the silicon probe electrode in the pyramidal layer for the sleep data in B. The population rate was computed as the average rate across all cells in 100 ms bins. (D) Cross-correlation between the population rate and silicon probe LFP power for different behavioral states. Top row shows results for theta LFP power and bottom row for slow gamma LFP power. Vertical red bars indicate locations of CA1 layers as in B. Same recording as in B. (E) Mean vector length of phase locking between spike times of single cells and LFP phase. Mean ± SEM (smaller than line width). Columns show the same conditions as in D. Line colors indicate different animals. Horizontal red bars indicate locations of CA1 layers as in B and D.

Figure 5B shows 30 s extracts of movement signals, spiking activity, slow gamma (25–50 Hz) and theta (6-10 Hz) band LFP signals for a sleep and subsequent open field recording for an example animal. During sleep, spiking activity and LFP signals across all layers showed strong co-fluctuations, possibly reflecting cortical up and down states (Battaglia et al., 2004). These co-fluctuations extended across all probe layers but were not present in the “Active” or the “Quiescent” open field conditions. To test if this is true for the whole recording, we computed the cross-correlation between the population spike rate (defined as the sum across all cells in 100 ms second bins) and the power in the different LFP channels (Figures 5C,D). Cross correlations extended across all silicon probe electrodes and were strongest close to the pyramidal layer from which the single cells were recorded (Figure 5D). In contrast, cross-correlations were largely absent in the “Active” state and confined to the stratum radiatum in the “Quiescent” state of the open field recording. This effect did not depend on LFP power as this was generally stronger during open field recordings and firing rates of single cells were also higher during the active state (Figure S4). Thus, on a population level, spiking activity and LFP power show distinct behavioral state-dependent patterns across different CA1 layers.

We further asked if the association of the firing of single spikes to the phase of LFP oscillations on a millisecond time scale (phase locking) might exert a similar depth dependence. In contrast to the population activity, there was discernible phase locking patterns across all layers and conditions (Figure 5E). The dependency on LFP probe channel was largely consistent between pre and post sleep conditions for the theta band (permutation tests; *p >* 0.5 for all mice), exhibiting a clear peak in the stratum radiatum, but differed significantly for slow gamma (permutation tests; *p <* 0.001 for all mice). However, the depth-dependence of theta band phase locking was significantly different between the active and sleep conditions (permutations tests; *p <* 0.001 for all conditions for each mouse; Bonferroni correction).

These results demonstrate that the Hybrid Drive enables a detailed characterization of the relationship between layer-specific and across-layer neural activity during different behaviors in freely moving mice.

### Across-layer information improves prediction accuracy of CA1 pyramidal layer cell responses

We next investigated how multiple behavioral variables and neural activity across different hippocampal layers contribute to the firing of single cells within the CA1 pyramidal layer. CA1 pyramidal cells show selectivity to different behavioral variables, including the animal’s position, head direction, and running speed (O’Keefe and Dostrovsky, 1971, McNaughton et al., 1983, Muller et al., 1994, Skaggs and McNaughton, 1998, Knierim, 2002). At the same time, CA1 cells can be coupled to internal network state, in particular by firing in relation to theta oscillations (Skaggs et al., 1996, Csicsvari et al., 1999, Mizuseki et al., 2011). To simultaneously characterize selectivity of single CA1 cells to multiple behavioral variables and theta activity, we adapted a generalized linear model (GLM) with Poisson response distribution (Figure 6A) (Burgess et al., 2005, Acharya et al., 2016, Hardcastle et al., 2017). A major advantage of a GLM-based approach over traditional tuning curves is its robustness to the interdependence of encoded variables which can lead to bias when analyzing the impact of these variables separately (Hardcastle et al., 2017, Stevenson, 2018, Laurens et al., 2019). For each cell, a GLM was trained to predict the cell’s firing in 100 ms bins using the animal’s position on the linear track, head direction, and running velocity (where positive/negative velocity corresponds to the animal running to the far/near end of the track). Additionally, we included neural variables derived from the chronic silicon probe in the Hybrid Drive: theta (6–10 Hz) LFP amplitude extracted from either the electrode in the pyramidal layer (in which the tetrodes were positioned) or from all CA1 electrodes (Figure 6A, bottom right). The latter allows for integration of across-layer information which cannot be captured by the tetrodes alone.

**Figure 6:**
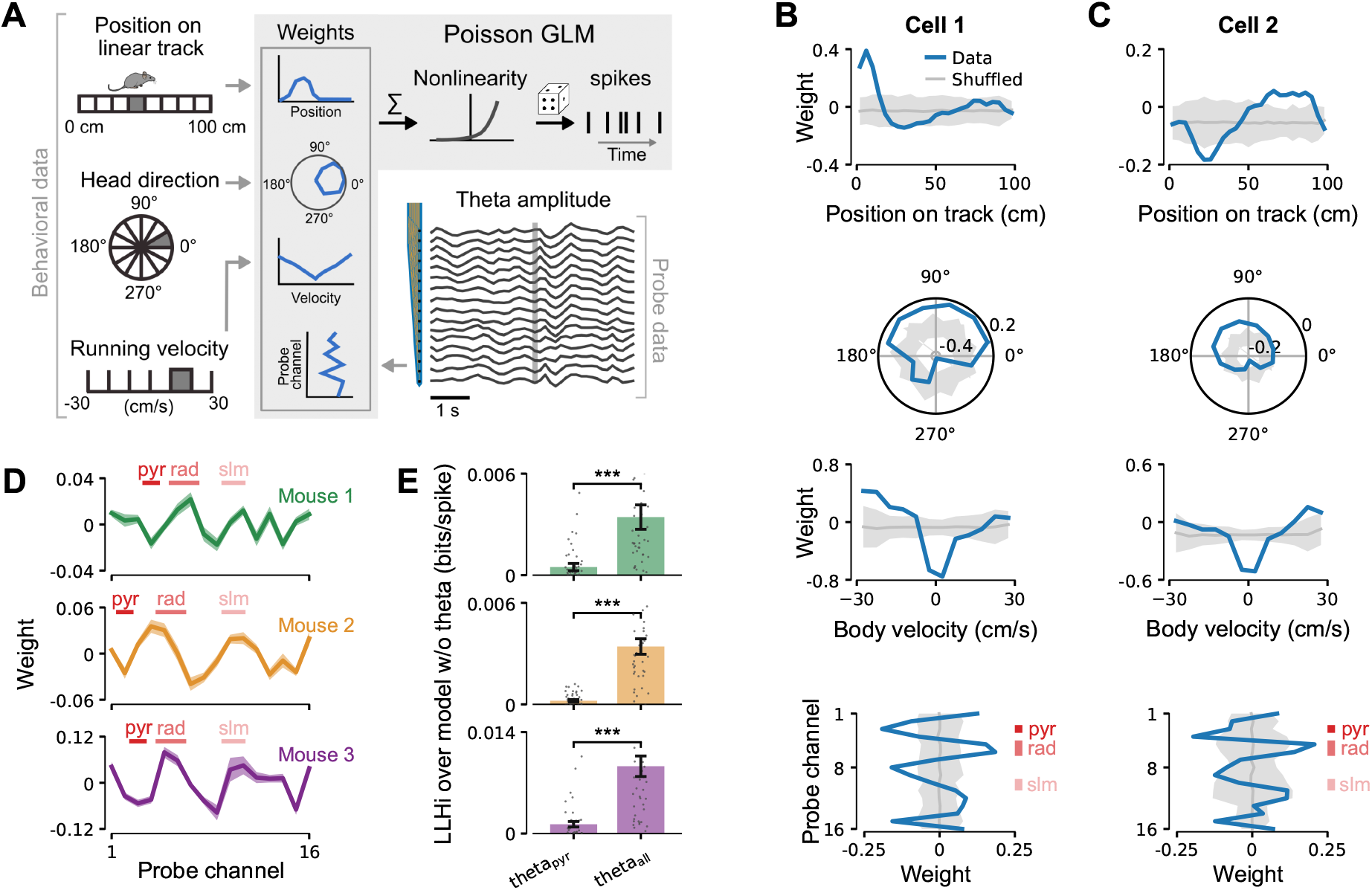
Across-layer information improves prediction accuracy of CA1 pyramidal layer cell responses. (A) Behavioral variables and silicon probe data were used to fit the parameters of a Poisson Generalized Linear Model (GLM) to predict the response of simultaneously recorded pyramidal layer cells. The gray bin indicates the current value. For probe data, the vertical gray line indicates the current theta band (6–10 Hz) LFP amplitudes. Position on track and running velocity bin counts reduced for illustration purposes. (B) Fitted GLM weights for an example CA1 pyramidal layer cell recorded using the tetrodes in the Hybrid Drive. Solid blue lines show weights of model fitted on the measured spike train. Gray line and shaded gray area show mean and 3 SD of shuffled spike train, respectively (*n* = 1000 shuffles). Vertical red bars in bottom plot indicate locations of pyramidal, radiatum, and slm layers relative to the silicon probe electrodes. (C) The same as in B for another example cell. (D) Summary of theta amplitude GLM weights for three animals. Horizontal red bars indicate locations of CA1 layers relative to the silicon probe electrodes as in B and C. Positive (negative) weights correspond to an increase (decrease) in single-cell firing and are consistent for pyramidal layer, radiatum and slm across mice. Mean ± SEM across cells for each animal (mouse 1: 51 cells, mouse 2: 47 cells, mouse 3: 41 cells). Only cells with a firing rate of at least 2 spikes/s were included in the analysis. (E) Prediction performance of a GLM that includes theta amplitudes from all silicon probe electrodes (theta_all_) compared to a GLM that only includes theta LFP amplitude for the electrode in the pyramidal layer (theta_pyr_). Cross-validated log-likelihood increase (LLHi) relative to a model without silicon probe data. Dots indicate single cells for the same data as in D. Wilcoxon signed-rank test on log-likelihood, ***p < 0.0001. Mean *±* SEM.

Many CA1 pyramidal layer cells showed tuning to multiple behavioral variables and theta amplitude at different depths (Figures 6B,C). To determine the significant GLM weights for each variable, we also computed weights for the same model (i.e. the same hyperparameters) after shuffling the spike train for each neuron; significant GLM weights were defined as those that exceeded 3 standard deviations of the shuffled weights (*n* = 1000 shuffles). Significant theta amplitude weights were generally large for silicon probe electrodes in the stratum pyramidal, stratum radiatum and stratum lacunosum moleculare (“pyr”, “rad”, and “slm” in Figures 6B,C). This observation was consistent across animals (Figure 6D). Importantly, inclusion of across-layer theta power significantly improved cross-validated log-likelihood compared to a GLM with only pyramidal layer theta amplitude (Figure 6E; Wilcoxon signed-rank test; mouse 1, *p* = 7.5 *·* 10^−7^; mouse 2, *p* = 1.1 *·* 10^−10^; mouse 3, *p* = 4.5 *·* 10^−12^).

We conclude that across-layer LFP signals provide additional information for single cells that is not captured by behavioral variables and the LFP signal within the pyramidal layer alone. Moreover, the consistent structure of theta GLM weights across mice support the idea that theta oscillations in the pyramidal, radiatum, and slm layers provide distinct afferent inputs to pyramidal layer cells (Schomburg et al., 2014, Lasztóczi and Klausberger, 2014).

## Discussion

Here we present the Hybrid Drive, a chronic implant device combining tetrodes (up to 14) and silicon probes (up to 2) in compact, flexible arrays (Figure 2, Figure S2). The drive is light enough to study a wide range of naturalistic behaviors; this includes standard paradigms such as running on a linear track (Figure 4) and open field exploration (Figure 5) but also interactions with objects in the environment (Figure 4) and, importantly, natural transition to sleep (Figure 5).

### Comparison with other extracellular recording devices

The Hybrid Drive matches the number of tetrodes of other state-of-the-art chronic implant devices (flex-Dive and shuttleDrive; Figure 7), combining at the same time up to two 16 channels silicon probes (for a total of 88 recording channels). The easy-to-build array allows the user to create a wide range of designs to tackle the target region of interest at best. The tetrodes and silicon probes can be part of the same array (Figure 2) or be divided into different arrays, simultaneously entering the brain at different angles (Figure S2). This greatly expands the amount of possible designs, and it’s not possible with simpler single-drive probe drives (Vandecasteele et al., 2012, Chung et al., 2017, van Daal et al., 2021) (but see Vöröslakos et al. (2021) for combining multiple drives).

**Figure 7:**
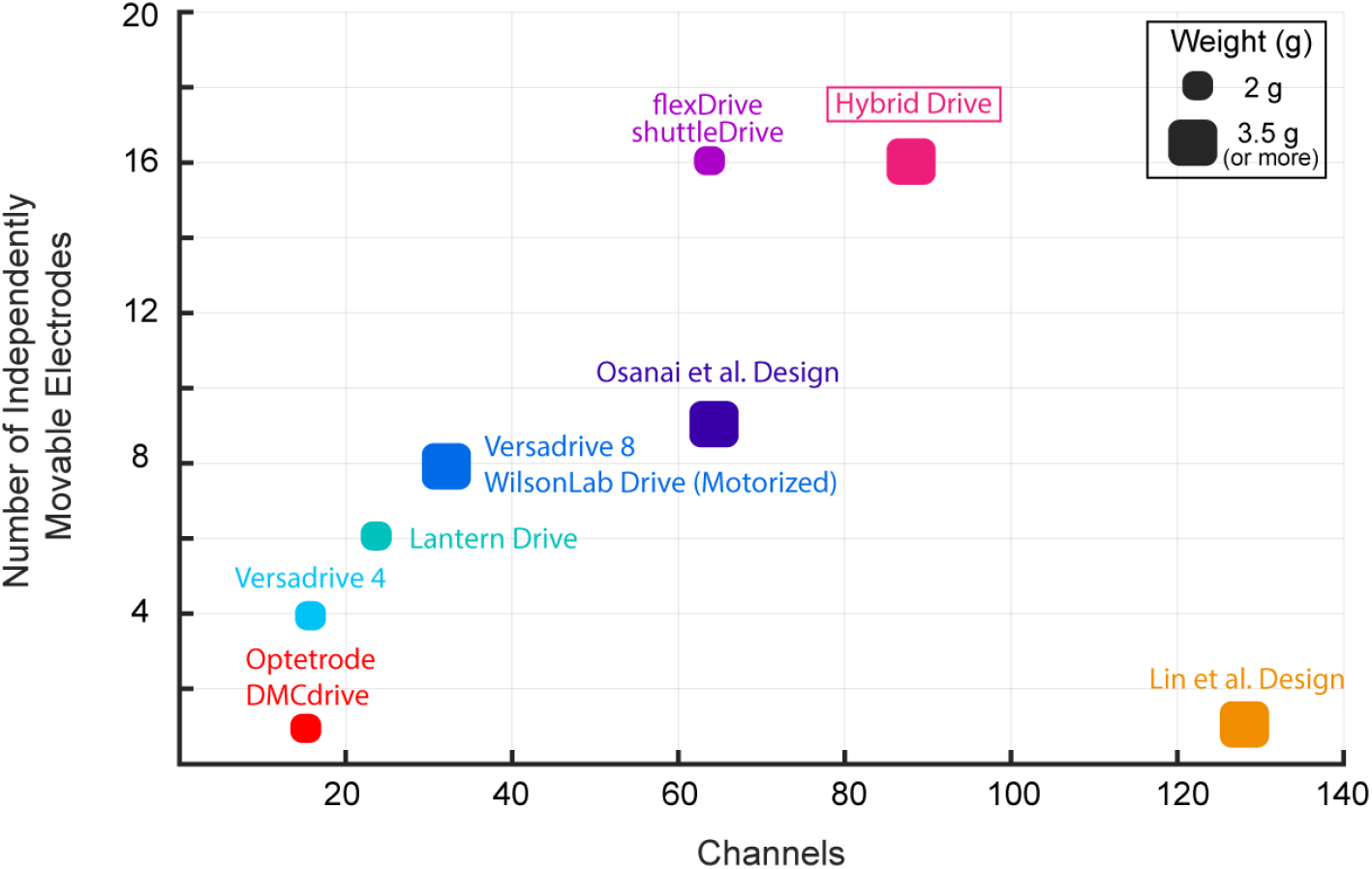
Comparison between tetrode implant devices for mice. Our system register the highest number of recording channels, while combining independently movable tetrodes and silicon probes in unique array designs (Lin et al., 2006, Yamamoto and Wilson, 2008, Battaglia et al., 2009, Kloosterman et al., 2009, Anikeeva et al., 2012, Voigts et al., 2013, Osanai et al., 2019, Voigts et al., 2020).

Osanai et al. (2019) developed another implant device combining silicon probes with tetrodes. However, our design provides two important advantages. First, their design can combine only nine tetrodes with a single shank 32-ch silicon probe. The two technologies can not be combined in the same array, limiting the range of the achievable designs. Second, the higher weight (5.4g) and larger size (large cone shape) of the their design might interfere with behavioural experiments in which the mouse has to actively interact with the environment (Figure 4) or with other conspecifics.

Parallel recordings of single units and LFP have been carried out using high-density silicon probes as well (Vandecasteele et al., 2012, Steinmetz et al., 2019, 2021). However, tetrodes still present some key advantages with respect to other methods for extracellular electrophysiological recordings. First, they provide a great recordings stability over time. Individual neurons can be consistently recorded through weeks and months (Fraser and Schwartz, 2012), or even a year (Voigts et al., 2013). This enables longitudinal experiments that follow learning processes. The same degree of durability of the ensemble recordings is not achievable using silicon probes (Karumbaiah et al., 2013, Kozai et al., 2015). Second, each individual tetrode can be independently lowered in the target area and precisely adjusted until the electrophysiological markers of the target layer are found. This is especially useful when positioning tetrodes in superficial structures (or cortical layers) that can be deformed or damaged by the probe insertion (during surgery). Last, our design allows to measure the LFP with a separate set of recording sites than those used to record single unit activity, even within the same area. This crucially avoids any spillovers of spiking activity into the LFP, a critical problem when using analytical approaches that explore LFP-single spike interactions (Buzsáki et al., 2012).

Our novel system offers great latitude in the placement of the tetrodes and the probes, fitting the unique geometry of individual brain areas (Figure 2, Figure S3) or the combinations of multiple areas (Figure S2). We can place a large number of recording sites in a thin layer (Figure 2), yielding large neuronal populations while recording the LFP from the same (or different) regions for an extended period of time (Figure 3). The current EIB is designed to incorporate two 16 channels linear silicon probes, optimized for the CA1/CA3 region of the mouse hippocampus. However, other 16 channels flexible-shaft silicon probes, with unique geometrical organization of the contacts, are readily available. Denser, non-linear organizations can be used to infer the sub-layer origin of the recorded units, e.g. deep vs superficial hippocampal pyramidal cells (E16R+R-20-S1-L20 NT; Mizuseki et al. (2011)), or shaft with electrodes organized in more spaced configuration that can sample LFP across multiple areas (E16+R-250-S1-L8 NT). This is not possible with simpler single-drive probe drives (Vandecasteele et al., 2012, Chung et al., 2017, van Daal et al., 2021) (but see Vöröslakos et al. (2021) for combining multiple drives).

Last, the current version of the Hybrid Drive requires flexible-shaft silicon probes (Herwik et al., 2011); changes in the drive design or the EIB layout might be required to fit less flexible silicon probes such as multi-shank or “Neuropixels” probes (Jun et al., 2017, Steinmetz et al., 2021). For example, using a steel spring with longer arms to reduce the bending angle or adapting a drive mechanism with straight drives (Voigts et al., 2020) could help to house a wider range of silicon probes.

### The Hybrid Drive enables advanced analytical approaches

The combined study of local field potentials and neural ensembles is behind many key advances in our mechanistic understanding of neural information processing. Laminar resolved recording (and Current Source Density (CSD) analysis) have been instrumental in dissecting contribution of different microscopic and mesoscopic scale circuits to neural function (Schomburg et al., 2014). On the other hand, recording of large neural ensembles (hundreds on neurons simultaneously) enabled a better understanding of neural processes behind higher cognitive functions (Peyrache et al., 2009). Linking the two analytical approaches can provide unique insights on how different input streams to a cortical module shapes the organization of neural populations. In this paper, we show how our Hybrid Drive may be used to precisely and reliably study the association of neural ensemble firing with layer-resolved oscillatory patterns in small rodents, focusing on the CA1 region of the hippocampus.

The hippocampus is one of the brain regions where the functional connectivity and the link to oscillatory phenomena are best known (Schomburg et al., 2014), but the same analysis approaches have been successfully applied to the neocortex as well (Kerkoerle et al., 2014). For example, in Figure 5 we show how the correlations between population firing and LFP generators at different layers is affected differently by changes in behavioral state, with only the stratum radiatum retaining coherence outside of the sleep state (characterized by low-frequency driven high coherence throughout the layer; Figure 5D). Similarly, the phase locking pattern to theta and slow gamma show layer and behavioral state dependent effects (Figure 5E). For example, a clear peak in the stratum radiatum emerges in the sleep phase, which is absent during wakefulness. Our analytical approach, facilitated by the type of data recorded with the Hybrid Drive, also shows that the availability of the layer-specific LFP enables a better characterization of the coding properties of single hippocampal cells. A Poisson GLM including layer resolved theta power has significantly higher predictive power for cell firing compared to behavioral variables typically used in spatial coding (Hardcastle et al., 2017), with a distinctive pattern of layer contributions emerging (Figure 6). Interestingly, when predicting single-neuron firing, GLM weights for theta power change sign multiple times along the dorsoventral axis, suggesting differential roles of generators in different layers in firing neurons. We believe that this approach and characterization of neural activity may help shedding light on neural coding properties, for example in approaches including “neural” and “external” predictor variables as in our analysis, helping understand and the role of certain inputs in shaping sub-populations of simultaneously recorded cells.

### The Hybrid Drive is modular, easy to build and open-source

The Hybrid Drive is open source and we provide all required design files along with detailed building instructions. Our innovations add value to the widely-used existing flexDrive design (Voigts et al., 2013), which should further promote its adoption. Just like the flexDrive, the system is modular and could be combined with technologies for optogenetic manipulation of neural activity or miniature head-mounted cameras to capture detailed behavioral variables, such as whisking or eye movements (Meyer et al., 2018). Together with recent advances in marker-less tracking of specific body parts (Mathis et al., 2018) and analytical tools to analyze spike trains of neurons across brain areas (Grossberger et al., 2018, Keeley et al., 2020), the Hybrid Drive helps to increase the precision and range of questions that can be addressed about the neural basis underlying natural behaviors in freely moving mice and other small laboratory animals. Our new design fits a gap in the existing technology and it may become an important tool in the system neuroscientist’s toolkit.

## Supporting information

Hybrid Drive building protocol

## Acknowledgments

We thank Jakob Voigts, Bruce McNaughton, Oscar Chadney and Jan Klee for their comments on the manuscript and the Battaglia lab for helpful discussions and comments during different stages of this work. This work was supported by the Radboud Excellence Initiative (A.F.M.), the European Union’s Horizon 2020 research and innovation program (MGate, grant agreement no. 765549; M.G. and F.P.B.), and the European Research Council (ERC) Advanced Grant “REPLAY-DMN” (grant agreement no. 833964; F.P.B.).

## Author contributions

Conceptualization: M.G., A.F.M., and F.P.B.; Investigation and Data Curation: M.G.; Methodology and Software: M.G, R.E., R.P., and A.F.M; Resources: A.A. and F.P.B.; Formal Analysis and Visualization, M.G., R.P., and A.F.M.; Writing – Original Draft: M.G. and A.F.M; Writing – Review & Editing: M.G., A.F.M., and F.P.B.; Supervision and Funding Acquisition: A.F.M. and F.P.B.

## Materials and Methods

### Ethics statement

In compliance with Dutch law and institutional regulations, all animal procedures were approved by the Central Commissie Dierproeven (CCD) and conducted in accordance with the Experiments on Animals Act (project number 2016–014 and protocol numbers 0021 and 0029).

### Animals

9 male C57BL6/J mice (Charles River) were used in this study. 4 animals were implanted with an Hybrid Drive in the CA1 area. Additionally, as a proof-of-principle, one animal received a dual area Hybrid Drive implant (parietal cortex and visual/para-hippocampal cortex) and another animal a Hybrid Drive implant over the subiculum. 3 animals were implanted with a flexDrive for the object exploration experiment. All animals received the implant between 12 and 16 weeks of age. After surgical implantation, mice were individually housed on a 12-h light-dark cycle and tested during the light period. Water and food were available *ad libitum*.

### Probe fabrication

The flexibility of the silicon probes is an important feature that makes the current design of the drive possible. The flexible polyimide probes used in this study have been designed and fabricated by ATLAS Neuroengineering BV (Leuven, Belgium) and used in other design for freely behaving rats (Michon et al., 2016). The fabrication is done on 4 inch silicon wafers using standardized fabrication techniques established using micro-electromechanical systems engineering. First, a stress-compensated passivation stack of silicon oxide and silicon nitride is deposited and acts as an electrical insulator between the probe metallization and the underlying bulk silicon substrate. Next, the metallization is sputter deposited and patterned using a lift-off process. A second passivation layer has been deposited which protects the metallization against the cerebral fluids in the brain tissue. The passivation layer stack is patterned by reactive ion etching to expose the electrode contacts along the probe shaft and the bonding pads on the probe base. The electrode metallization layer is deposited and patterned using a lift-off technique, resulting in electrodes with surface area of 225*μ*m^2^ made of iridium oxide. Following the definition of the electrodes, the passivation and insulation layers are patterned using reactive ion etching to define the probe outline and exposing the build silicon substrate. Next, trenches with vertical sidewalls are etched into the bulk silicon substrate using deep reactive ion etching followed by a wafer thinning process to end up with 50*μ*m thin silicon probes. Finally, the probes are assembled with the highly flexible PI ribbon cable using a flip-chip bonding process.

### Hybrid Drive Assembly

All parts and steps required to build a Hybrid Drive are summarized in a separate step-by-step protocol (https://github.com/MatteoGuardamagna/Hybrid_drive). Briefly, the drive mechanism of the Hybrid Drive is based on an existing design (flexDrive, Voigts et al. (2013)). Tetrodes were made from HM-L coated 90% platinum/10% iridium 12.5 *μ*m diameter wire (California Fine Wire, USA). The guide arrays containing the tetrodes were assembled by building up layers of polyimide tubes, according to the target areas, and fixing them with epoxy resin (Araldite Rapid, Araldite, UK). For the silicon probe used in this study (E16R_60_S1_L20_NT with 60 μm pitch between contacts, ATLAS Neuroengineering, Belgium), we utilized a 33ga polymide tubing for the entire structure supporting the shaft instead of a combination of 33ga and 38ga polymide tubing used to build the tetrode slots. Depending of the width of the shaft and the organization of the guide array a larger diameter size of polymide tube (e.g. 26ga) can be chosen for the silicon probe. We opted to keep the same size to be able to fit all 14 tetrodes and the silicon probe in the CA1 region. For the CA1 and CA1/subiculum design we used a 5×3 array design to cover the dorsal CA1 area along its proximo-distal axis (Figure 2A) For the dual area implant design two bundles were used. A 4×3 bundle was positioned over the parietal cortex and a single tube was used to position the probe in the visual/para-hippocampal cortex (Figure S2). The spring-screw mechanism of the flexDrive allowed precise positioning and re-positioning of individual tetrodes (Figures S1B,C). This mechanism has been adapted and implemented in the Hybrid Drive to allow for independently movable silicon probes (Figure S1D). First, the portion of polyimide tubing that will create the upper guide tube must be slid along the silicon probe shaft. The length difference between the upper and lower guide tube determines the total travel distance of the probe (the maximum recommended distance is 2 mm; Figures S1A,D). After positioning each tetrode guide tube in the slots of the drive body, the silicon probes were inserted in the corresponding lower guide tubes. Finally, The EIB was glued to the drive body using epoxy resin and the silicon probes’ ZIF plugs were inserted into the ZIF connectors on the EIB. The current version of the EIB (Figure 1D) contains two 16 channel ZIF connectors. The flexible cables of the silicon probes were protected using lightweight cone shields attached to opposite ends of the EIB (Figure 1A,B).

### Surgical Procedures

All surgeries were performed on experimentally-naive mice. Mice were anaesthetized with 1.25% – 2% isoflurane and injected with analgesia (Carprofen, 5 mg/kg SC). Mice were placed in a stereotaxic frame and antibacterial ophthalmic ointment (Alcon, UK) was applied to the eyes. A circular piece of dorsal scalp was removed and the underlying skull was cleaned and dried. 3 stainless steel M0.8 screws were used to secure the drive (1 ground screw in the frontal plate, 1 screw in the parietal plate opposite to micro-drive, 1 screw in the occipital plate). For the CA1 implant, a craniotomy was made over the right cortex (top-left corner at AP: −1.20 mm; ML: 0.6 mm relative to bregma; bottom-right corner at AP: −2.30 mm; ML: 2.10 mm relative to bregma) using a 0.9 burr drill. The dura was removed and the array of the drive was slowly lowered into the brain with the silicon probe shaft already adjusted at the final depth (*≈* 2.00 mm; Figure 2A). The guide tube array was lowered right above the brain surface and the craniomoty was filled with sterile vaseline to protect the brain and the array from cement flowing in. The drive was cemented onto the skull using dental adhesive (Superbond C&B, Sun Medical, Japan) and tetrodes were individually lowered into the brain (5 turns - *≈* 900*μ*m) using the screw-spring mechanism. Mice were allowed to recover from surgery for at least seven days before experiments began. The same procedure was carried out for the subiculum and dual area implant. For the subiculum a craniotomy was made over the right cortex (top-left corner at AP: −2.40 mm; ML: 1.32 mm relative to bregma; bottom-right corner at AP: −3.20 mm; ML: 2.20 mm relative to bregma; using a 0.9 burr drill.) Two craniotomies were done over the right cortex for the dual area implant. For the anterior bundle: top-left corner at AP: −1.46 mm; ML: 1.00 mm relative to bregma; bottom-right corner at AP: −2.30 mm; ML: 1.50 mm relative to bregma; using a 0.9 burr drill. For the posterior bundle: AP: −4.75 mm; ML: 3.25 mm relative to bregm; using a 0.5 burr drill.

### Neural and behavioral data collection

From post-surgery day 3 onward, animals were brought to the recording room and electrophysiological signals were investigated during a rest session in the home cage. Each day tetrodes were individually lowered in 45/60*μ*m steps (1/4 of a screw turn) until common physiological markers for the hippocampus were discernible (SWR complexes during sleep or theta during locomotion). The majority of the tetrodes reached the target location (CA1 pyramidal layer) in 7-10 days. Silicon probe signals were used as additional depth markers. Electrophysiological data were recorded with an Open Ephys acquisition board (Siegle et al., 2017). Signals were referenced to ground, filtered between 1 and 7500 Hz, multiplexed, and digitized at 30 kHz on the headstages (RHD2132, Intan Technologies, USA). Digital signals were transmitted over two custom 12-wire cables (CZ 1187, Cooner Wire, USA) that were counter-balanced with a custom system of pulleys and weights. Waveform extraction and automatic clustering were performed using Dataman (https://github.com/wonkoderverstaendige/dataman) and Klustakwik (Rossant et al., 2016), respectively. Putative excitatory pyramidal cells were discriminated using their auto-correlograms, firing rates and waveform information. Specifically, pyramidal cells were classified as such if they had a mean firing rate < 8 Hz and the average first moment of the autocorrelogram (i.e., the mean value) before 8 msec (Csicsvari et al., 1999). Clustered units were verified manually using the MClust toolbox.

During all experiments, video data was recorded using a CMOS video camera (Flea3 FL3-U3-13S2C-CS, Point Grey Research, Canada; 30 Hz frame rate) mounted above the arena. Animal position data was extracted offline using a convolutional deep neural network trained using transfer learning (Mathis et al., 2018).

### Behavioral paradigms

We used three different behavioral paradigms: running on a linear track, open field exploration, and object exploration. In the linear track experiment (Figures 4A–D; Figure 6) mice were positioned at one end of track of a 1-meter long track with the task to run to the other end to collect a reward (a piece of Weetos chocolate cereal). After the animal consumed the reward, another reward was positioned on the opposite end. A lap was defined as an end-to-end run where the animal’s body started from the first 10 cm of the track and reached the last 10 cm at the other end of the track without returning to its initial position. Recordings typically lasted between 20 and 30 minutes and experiments were performed on 10 consecutive days.

In the open field experiment (Figure 5), animals were free to explore an unfamiliar square arena (45 cm x 45 cm) for about 30 minutes. This was preceded and followed by a rest session in the animal’s home cage (“Pre sleep” and “Post sleep”) that typically lasted 60 minutes each.

The object exploration experiment (Figures 4F–J) was adapted from Deshmukh and Knierim (2011). Briefly, animals were free to explore a 75 cm x 75 cm square arena containing 3 objects for 5–7 consecutive trials (10 minutes each). Between trials, mice were returned to their home cages for 5 minutes. In each animal, the same procedure was performed on at least 3 consecutive days, resulting in 18 trials per animal that were included in the analysis. Object locations were kept constant for each animals throughout all trials.

### Histology

After the final recording day tetrodes were not moved. Animals were administered an overdose of pentobarbital (300 mg/ml) before being transcardially perfused with 0.9% saline, followed by 4% paraformaldehyde solution. Brains were extracted and stored in 4% paraformaldehyde for 24 hours. Then, brains were transferred into 30% sucrose solution until sinking. Finally, brains were quickly frozen, cut into coronal sections with a cryostat (30 microns), mounted on glass slides and stained with cresyl violet. The location of the tetrode tips was confirmed from stained sections.

### Neural data analysis

For each identified spike unit, the signal-to-noise ratio (SNR) was used as a measure of electrophysiological signal quality (Figures 3A–E). The SNR was computed as the ratio between the peak absolute amplitude of the average spike waveform and noise on the tetrode channel on which the peak amplitude occurred (Meyer et al., 2018, Magland et al., 2020). To compute separate SNRs for “Active” and “Quiescent” conditions for each unit (Figures 3D,E), recordings were first segmented using the 3-axis accelerometer signals from one of the recording headstages (sampled at 1000 Hz). Calibration of the accelerometer and extraction of head movement signals followed previous work (Meyer et al., 2018). Time points at which accelerometer-derived head movement signals exceeded 0.02 m/s2 were assigned to the “Active” condition whereas all other time points were assigned to the “Quiescent” condition.

To assess the recording quality of the chronically implanted silicon probe across days, we analyzed the properties of sharp wave ripple (SWR) events (Figures 3G–K). LFP signals were down-sampled to 1 kHz and the analysis was restricted to periods when the animal was in the “Quiescent” state. First, continuous data within these periods were filtered between 100-250 Hz using the “eegfilt” function (Delorme and Makeig, 2004). Then, for each animal, the channel with the highest ripple amplitude was selected and defined as the pyramidal layer channel. All subsequent analysis steps were based on this channel. The envelope of the filtered signal was estimated in 10 ms windows, representing the ripple amplitude of each time series. Ripples were detected using a threshold (mean of the signal + 3 std) on the envelope. Additionally, the duration of the event had to exceed 40 ms. SWR duration was calculated using onset and offset of each event by identifying the points that crossed the mean of the signal + 1 std, before and after the detected peak in the envelope. SWR power was defined as the power at the peak of the envelope for each individual ripple event (Figure 3H). SWR duration was defined as the interval between the onset and the offset of each ripple event (Figure 3I). SWR frequency was calculated as the frequency at the peak value of the spectrogram computed for the SWR event (Figure 3J). The proportion of SWR was defined as the total number of SWRs divided by the duration of the quiescent state (Figure 3K). CA1 layers were estimated for each animal by combining the information provided by the depth profile of SWR complexes (Figure 3G), the theta power depth profile (Figure S4A), and the distance and displacement of the silicon probe contacts. Theta and sow gamma oscillation have been extracted filtering the continuous data from the silicon probe using the “eegfilt” function (Delorme and Makeig, 2004) (6-10 HZ and 20-50 Hz respectively).

### Generalized-Linear Model (GLM) with Poisson observations

We adopted a Poisson GLM to investigate how multiple behavioral variables and across-layer LFP signals from the silicon probe contribute to single-cell firing in the linear track experiment. The implementation of the GLM largely followed that of Hardcastle et al. (2017). Briefly, spike trains of single cells were binned at 100 ms. The animal’s head position, head direction, and running speed were extracted from video data using a convolutional neural network (Mathis et al., 2018). Linear interpolation was used to temporally align the extracted behavioral data to the bin centers. For the position data, the linear track was divided into 25 non-overlapping bins (bin width 4 cm) along its major axis. For each time step, the indicator function (either 0 or 1) was used to indicate if a spatial bin is occupied by the animal. The same approach was used for head direction (24 bins of size 15°) and running velocity (12 bins of size 5 cm/s). As the firing of place cells can depend on the animal’s running direction on a linear track, we used running velocity rather than running speed (where positive/negative velocity corresponds to the animal running to the far/near end of the track). In addition to the behavioral variables, theta (6–10 Hz) LFP amplitudes were extracted from either the electrode in the pyramidal layer (in which the tetrodes were positioned) or from all CA1 electrodes were included.

For each cell, a separate Poisson GLM was fit to the data using gradient descent on the negative log-likelihood (using the “minimize” function in the scipy Python package; Oliphant (2007)). Additionally, roughness penalties were introduced to enforce smoothness of the position, head direction, and running velocity GLM weights (Hardcastle et al., 2017); an L2 penalty was used for the theta amplitude weights. The hyperparameters determining the strength of the roughness and L2 penalties were found for each cell using grid search cross-validation (*N* = 4 folds; 10 log-spaced values in the interval [2^−3^, 2^12^] for each hyperparameter). As the evaluation of all possible hyperparameter combinations is computationally expensive, we used Bayesian hyperparameter optimization (using “BayesSearchCV” in the scikit-optimize package with 100 iterations; Head et al. (2020)). For each cell, the prediction accuracy of the different models (theta_pyr_ and theta_all_ in Figure 6E) was assessed using nested cross-validation (*N* = 4 folds). That is, the hyperparameters were determined using cross-validation on the training data and predictions were performed on the validation data. The data in Figure 6E were computed using the Poisson log-likelihood as described in (Hardcastle et al., 2017). Using the Pearson correlation between measured and predicted spike trains yielded qualitatively similar results (Wilcoxon signed-rank test; mouse 1, *p* = 2.2 *·* 10^−5^; mouse 2, *p* = 2.7 *·* 10^−10^; mouse 3, *p* = 2.2 *·* 10^−8^).

## Supplementary Figures

**Figure S1:**
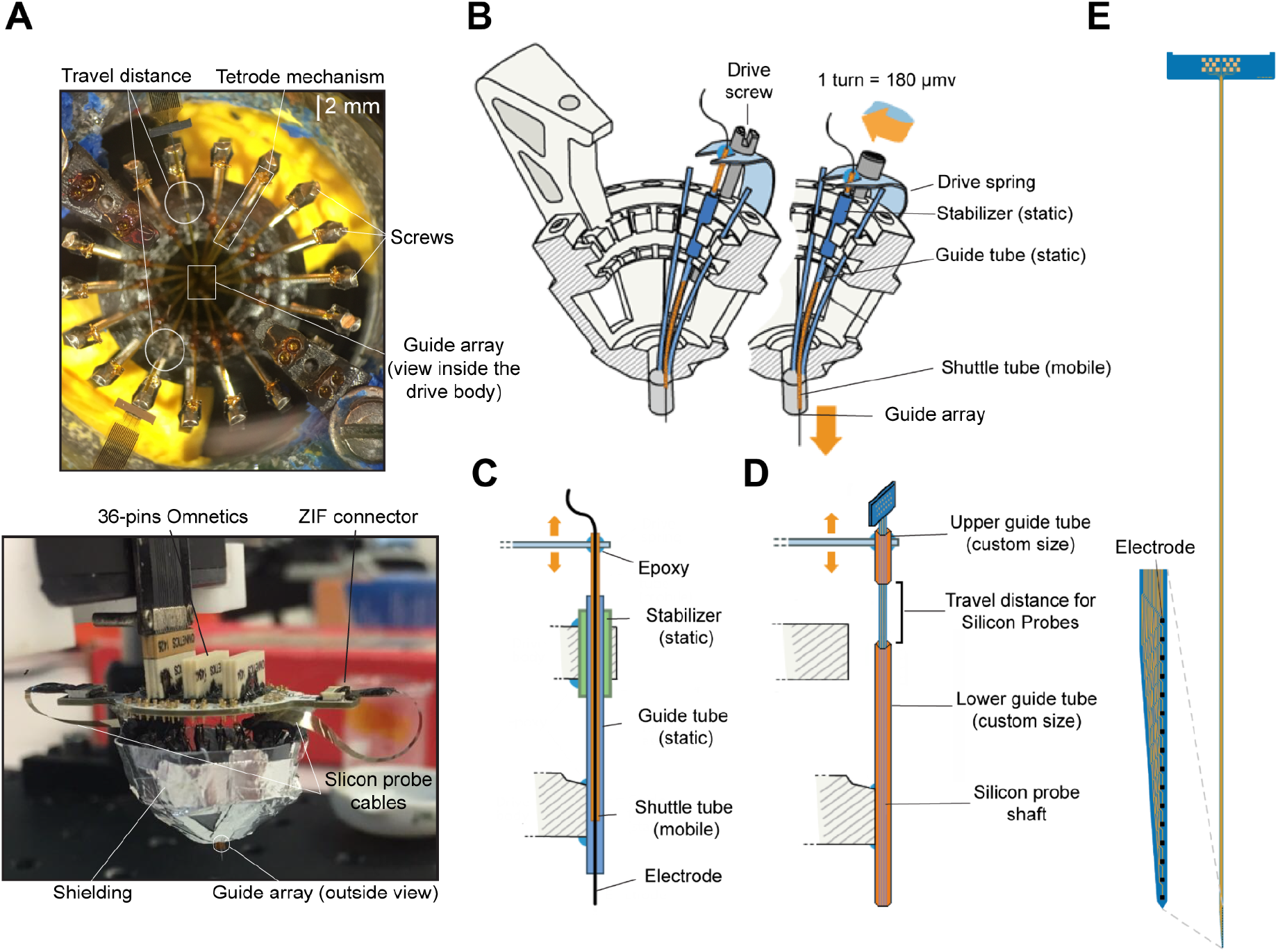
Silicon probe and tetrode mechanism. (A) The upper picture illustrate a top view of the drive body in which two silicon probes are positioned and the spring-screw mechanisms for tetrodes are assembled as well. The travel distance for the silicon probe is highlighted in the white circle. Lower picture illustrate the assembled drive, before the lateral cone shielding is applied. (B) Illustration of the spring-screw mechanism and the drive body, adapted from Voigts et al. (2013). (C) Detailed illustration of the spring-screw mechanism ensuring the independent movement of the tetrodes, adapted from Voigts et al. (2013). (D) Detailed illustration of the adapted spring-screw mechanism ensuring the independent movement of the silicon probes. (E) Illustration of the silicon probe used for the CA1 design.

**Figure S2:**
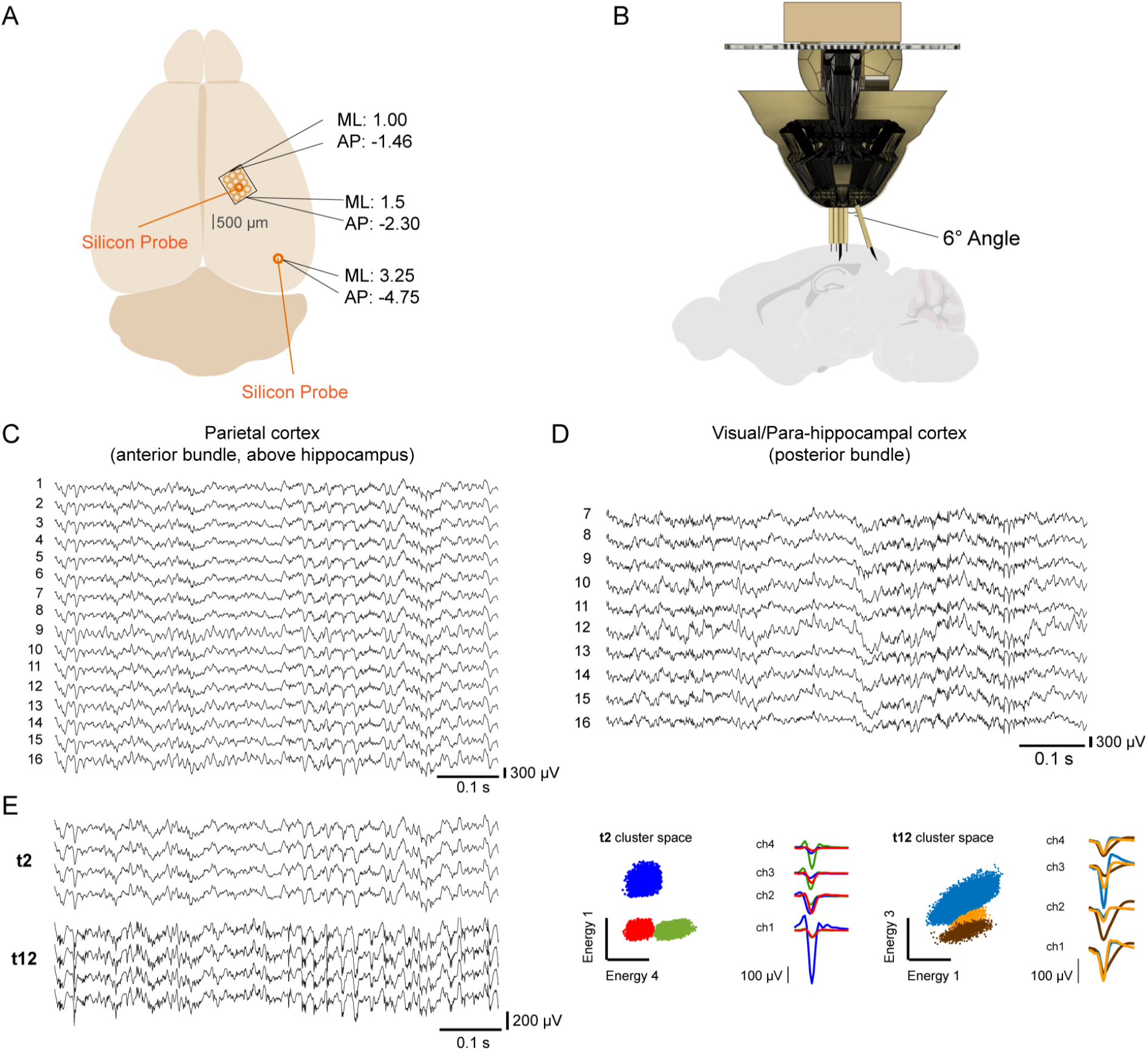
Dual area design with the Hybrid Drive. (A) Representative illustration with the coordinates of the two craniotomies. Brain illustration adapted from SciDraw (doi.org/10.5281/zenodo.3925941). (B) Representative illustration of the bundle organization of the dual area implant. The posterior bundle is glued at a 6° angle, with respect to the anterior bundle. Sagittal brain illustration adapted from SciDraw (doi.org/10.5281/zenodo.3925911). (C) Broadband signal (LFP) of the silicon probe in the parietal cortex above the hippocampus (anterior bundle). (D) Broadband signal (LFP) of the silicon probe in the visual/para-hippocampal cortex (posterior bundle). Only 10 contacts are reported because the probe was not fully lowered in the brain. The portion of the shank containing the first 6 contacts was left inside the guide tube array. Same time frame as in (C). (E) Left, broadband signal (LFP) from two example tetrodes (t2,t12 - four channels each), positioned in the parietal cortex over the hippocampus. t2 was left more superficial (2 screw turns, ≈ 360 *μ*m). t12 was driven deeper (4 screw turns, ≈ 720 *μ*m). Same timeframe of (C) and (D). Right, representative spike recording from the two tetrodes. 2D cluster projection of the recorded spikes and corresponding average spike waveforms.

**Figure S3:**
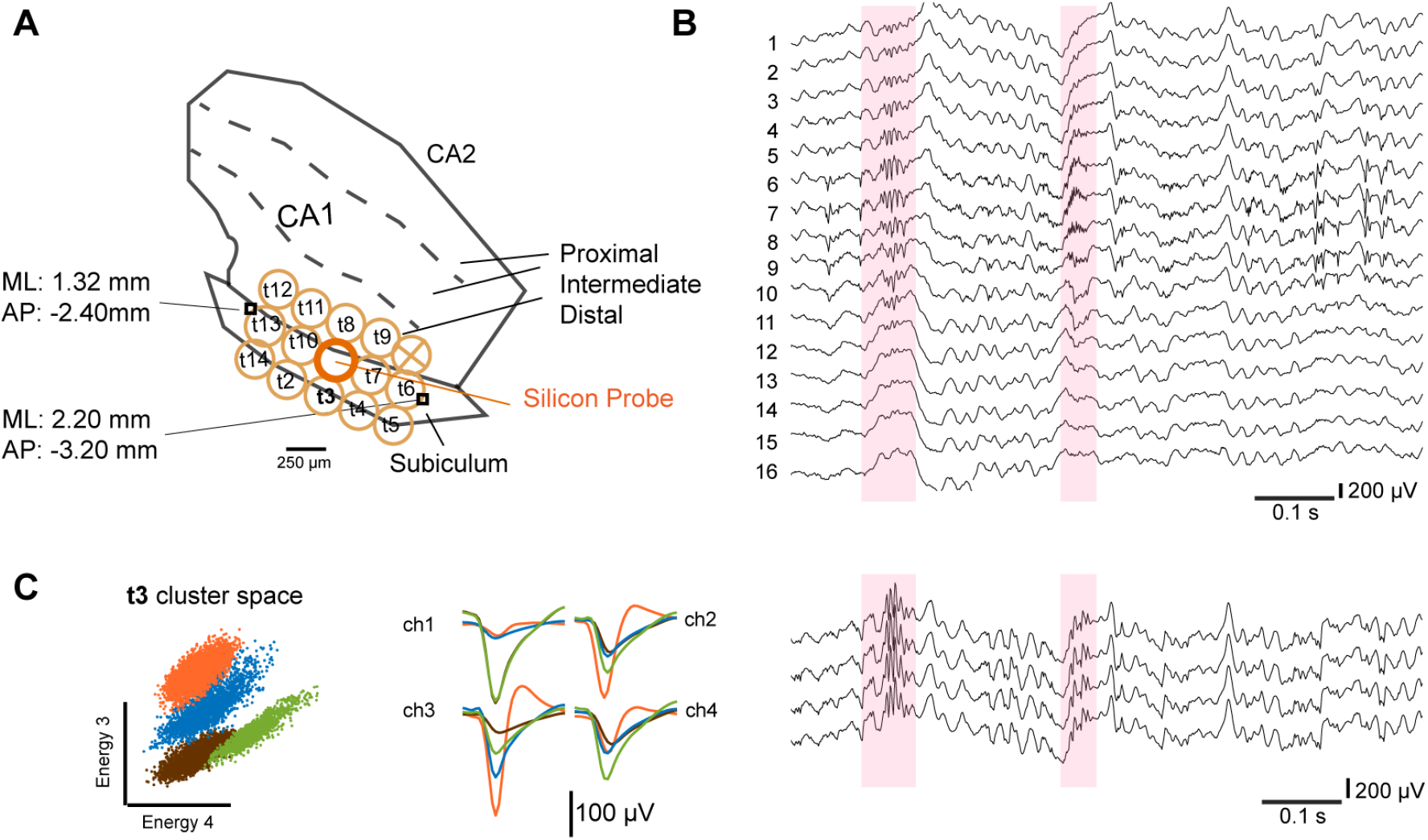
Array design and recordings from other hippocampal sub-regions. (A) Coordinates of the craniotomy over the subiculum, with respect to CA1. The guide tube array is represented in orange, with the mapping of individual tetrodes (and silicon probe) over the target structures. (B) Broadband signal (LFP) of the silicon probe. Ripple example highlighted in light pink. Note how the ripple spans over multiple contacts (5 contacts, ≈ 300 *μ*m - in line with the thickness of the subiculum along those implant coordinates) and it’s missing the typical CA1 sharp-wave component (Figure 3F for reference). (C) Left, representative spike recording from tetrode 3 (t3). 2D cluster projection (Energy 3 and 4) of the recorded spikes and corresponding average spike waveforms. Right, broadband signal (LFP) from one example tetrode (t3 - 4 channels). Same timeframe as in (B).

**Figure S4:**
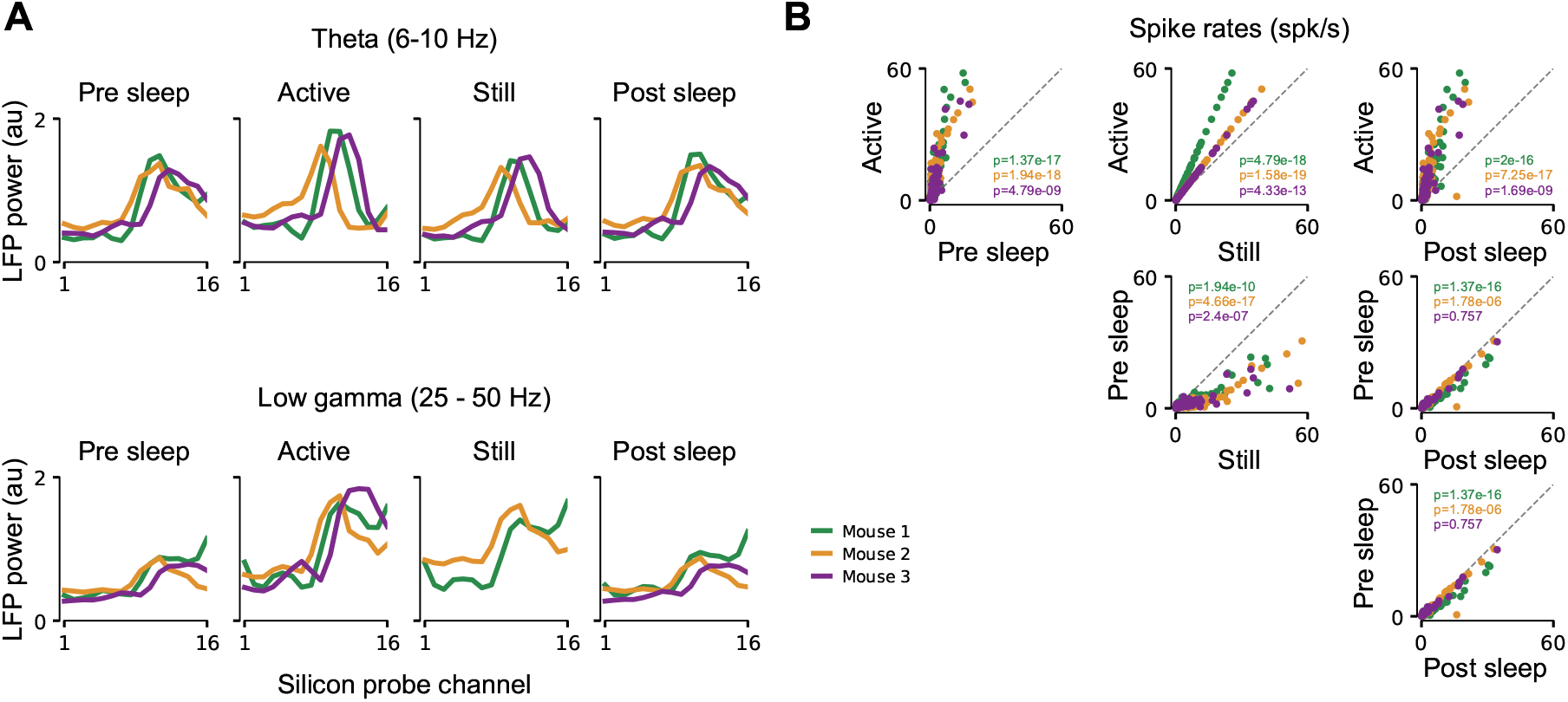
Comparison of across-layer LFP power and spike rates for open field exploration and sleep. Related to Figure 5. (A) LFP power for the different silicon probe electrodes spanning the vertical CA1 axis. Top: Theta band power for the different behavioral conditions in Figure 5. Bottom: The same but for the slow gamma frequency band. Colored lines indicate different animals. For each animal theta or slow gamma power in all conditions was normalized by the mean power in the active condition. (B) Average spike rates for single cells simultaneously recorded in the CA1 pyramidal layer. Same data and color scheme as in A. P-values were computed using a Wilcoxon rank-sum test (Bonferroni correction per animal).

