## Supplementary material for "The Hybrid Drive: a chronic implant device combining tetrode arrays with silicon probes for layer-resolved ensemble electrophysiology in freely moving mice": Hybrid Drive building protocol

### 0 Outline

This is a compendium on how to build the Hybrid Drive, a chronic implant device combining silicon probes and high density tetrode arrays. More information are found on the Hybrid Drive Github repository: `https://github.com/MatteoGuardamagna/Hybrid_drive` (Some parts of the Hybrid Drive are based on another implant device for extracellular physiology: the flexDrive. Check this Wiki for more detailed information on the flexDrive: `https://open-ephys.atlassian.net/wiki/spaces/OEW/pages/491529/flexDrive`.)

### 1 Drive Building

#### 1.1 Check/adjust the body of the drive

The drive body design is the in use for the the flexDrive:
`https://open-ephys.atlassian.net/wiki/spaces/OEW/pages/491578/Drive+body`. Few adjustment should be made to ensure a good organization of the guide array and the probe mechanism.

- Make sure screw holes are clear. If not, use a hand drill (0.45-0-55 diameter) to clear the opening.
  **Note:** The angle of the screw relative to the drive body is key for the tetrode adjustment (spring-screw) mechanism.
- Make sure the holes in the top part of the body are large enough for a screw or piece of tubing to slot in and secure the EIB to the body (at a later stage, see 2.6).
- Clear and/or enlarge guide tube grooves using a scalpel, to ensure that the stabilizer tube fits (see 2.9).
  **Note:** If the shape of the holders is damaged in this process, use superglue + accelerator to stick the stabilizer in position.
- Enlarge the bottom opening of the body using a dremel mounted with a drill tip (0.5 or 0.9 mm). It's important that the opening is capable of fitting the bundle, especially if you have it at and angle or more than one bundle. For CA1 implants, create a diagonal opening (shifted 30-40 degrees from the arms of the body to match the shape of the hippocampus).

**Note:** Make sure the diagonal is in line with the desired implant position throughout the process of building the drive, especially for the step 1.3.

- Place screws and lower them until the remaining visible threaded portion of the screw is approximately 3 mm.

### 1.2 Building the bundle tube array - specific 5x3 design

The version described here is specific for creating two 3x5 array.

- Place kapton tape in the shape of an "H" so that the sticky side of the middle bar if facing upwards. This is used to organize your tubing and make them stick correctly while you build up your array.
- Cut 15 pieces of tubing of approximately 6 cm. Choose the size of the silicon probe tubing depending on the organization of your guide array. Recommended sizes are 33ga (same as tetrode tubing) or 26ga.
- Lay down each tube segment so that the middle is centred on the tape. Align a row of 5 pieces of tubing on the tape without any gaps in between each other.
  **Note:** Whilst manipulating the tubing, always be careful not to squash or kink it. Hold gently by the extremity (the portion that will be cut off later on).
- Apply a thin layer of epoxy over the row of tubing, spread a few millimeters in either direction from the center.
- Repeat the process twice, adding a layer of 5 tubes each time.
- Let it dry for at least 1 hour.
- Cut the bundle in the middle with a decisive movement using a razor blade.
- Remove the bundle from the tape. Place a pair of fine forceps between tape and tubing and gently lift whilst sliding away from the middle.
  **Note:** Bundle should be approximately 2.2-2.5 cm in length.

### 1.3 Glue bundle to the drive body

Important: the position of the the array in the drive body is fundamental. Make sure the angle and orientation of the bundle array is correct and matches your coordinate planning.

- Place kapton tape in the shape of an "H" so that the sticky face of the middle bar is facing upwards. This is used to place and hold the bundle array in position while it is glued to the drive body.

- Place the drive body upside down in the appropriate direction, using tack on the body arms to hold it in place.
- Place the tube array in the body opening, aligned with the diagonal opening. The tube array should be sticking out by approximately 0.4 cm.

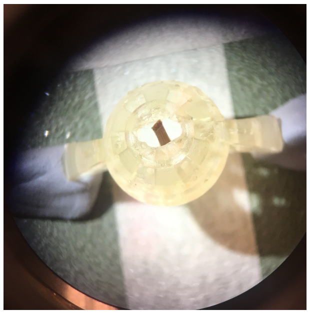

Figure 1: Drive body and tube array close up.

**Note:** It is best to have too much bundle than too little.

- Glue the tube array to the bottom of the drive in 3 steps using epoxy:
    1. Apply epoxy around the base, make sure the tube array is aligned as desired, the position can be adjusted as the glue dries. Leave to dry properly.
    2. Apply second layer of epoxy to reinforce the structure.
    3. Apply third layer at a later stage, when positioning the spring (see 1.4).

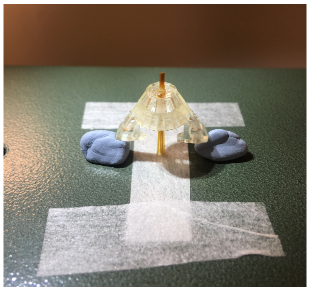

Figure 2: Tube array glued to the drive body.

#### 1.4 Assembling the spring

The spring design is the same used for the flexDrive:
`https://open-ephys.atlassian.net/wiki/spaces/OEW/pages/19988482/Drive+Spring`

- Use forceps to align, clip and hold in the place the extremities of the spring.
  **Note:** Make sure the shape of the spring is uniformly circular.

- Solder at the connection point using a minimal amount of solder.

#### 1.5 Add spring to the body

- Place the drive body upside down, using tack on the arms to hold it in place.

- Place the spring so that the bottom and sides align respectively with the bottom and arms of the drive body.
  **Note:** Indents on body made to hold the spring rarely fit. In fact, they might need to be removed for the spring to sit nicely on the drive. Can also try to tape the spring in place, at the level of the arms, before gluing.

- Glue the spring to the body using epoxy. Glue all around the bottom opening and add a drop at the level of each arm.

- Make sure the spring and drive body are aligned as the glue dries. The position can be adjusted as the glue dries.

- Let it dry for at least 1 hour.

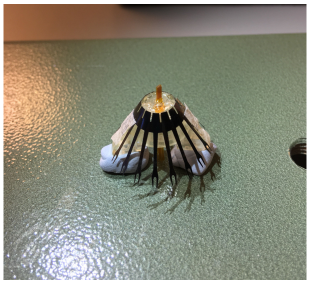

Figure 3: Drive body and spring.

#### 1.6 Place and organize guide tubes in the body

- Place the drive in its holder. The custom-made holder in the pictures is built from a plastic neck of a soda bottle. Portions of the bottleneck are

carved out. Those left hold the drive from the upper arms. Position the washers above each arm and tighten until the drive is stable. For extra stability, add a tiny drop of superglue between the washer of the holder and the arm of the drive body on either side.
**Note:** Center the drive as much as possible in the holder, generally be aware of the position of the guide tube array to avoid touching the side of the holder and causing damage.

- Identify the guide tubes that will be used for the probe. To help distinguish them from the tubes that will be used for tetrodes, insert a tube segment (38 ga) in the desired position from the bottom up.
  **Note:** At this point you can clean the bottom of the array, using gently a cotton bud + ethanol (very small amount).

- Place stabilizers:
  This step is only for guide tubes that will hold tetrodes.

  1. Cut a piece of stabilizer tubing (26 ga) of approximately 2 mm.
  2. Place the stabilizer over the guide tube.
  3. Using a pair of forceps, gently lower the stabilizer down the guide tube and lock it in the drive body (at the chosen position).
     **Note:** Pay attention to the organization of your guide tubes. Make sure that they are properly organized, not bent and mapped.
  4. Once all of the desired stabilizers have been locked to the drive, place a small drop of epoxy to secure the structure (make sure the epoxy enters inside the stabilizer).
     **Note:** Pay attention to not get any epoxy on the probe (or optic fiber) tubes that have not been positioned in the drive yet.
  5. At this point all guide tubes should be in position aside the guide tube of the silicon probe (and/or optic fiber).

- Cut the guide tube just above the stabilizer.

- To ensure better stability of the shuttle tube and the spring-screw mechanism build a bridge across the spring fork.

  1. Cut a piece of tube (33 ga) just larger than the width of the spring fork.
  2. Place a drop of superglue on the tube and position it on the spring fork.
  3. Add a drop of epoxy for stability.

- Place shuttle tube (38 ga).

  1. Cut a piece of shuttle tube (38 ga) for each of your guide tubes.

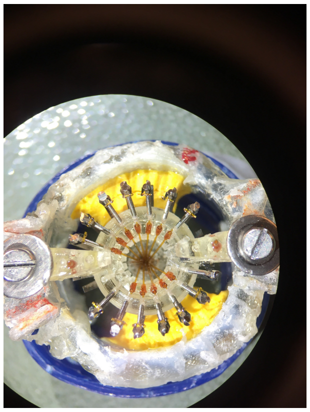

Figure 4: Guide tubes and stabilizers positioned in the drive body.

2. Insert the shuttle tube into the guide tube, locked into the drive body thanks to the stabilizer.
   **Note:** Pay attention to how much you insert the shuttle tube inside the guide tube. As a rule of thumb your shuttle tube should be visible (inside the guide tube) for at least 1 mm past the stabilizer.

- Epoxy shuttle tube to spring (and bridge).

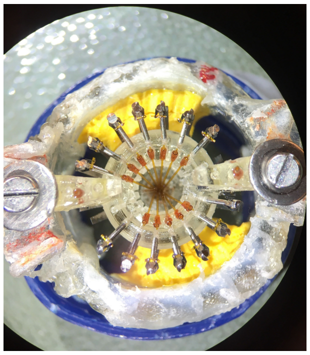

Figure 5: Inner tubes inserted in the guide tubes.

### 1.7 Prepare EIB

The custom EIB is made by ATLAS neuroengineering (https://www.atlasneuro.com/). The current design was studied to include 14 tetrodes and 2 silicon probes. The design of the EIB itself can be adapted, depending on the experimental needs.

- Solder wire from and to reference holes on bottom of the EIB.
  **Note:** Be careful not to overheat the EIB or damage the connections on

the board.

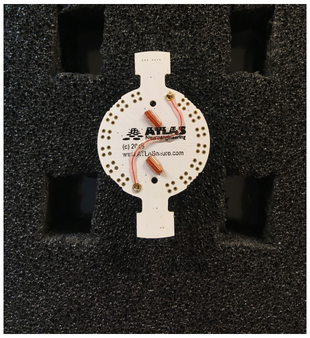

Figure 6: ATLAS EIB with ground connections soldered together.

#### 1.8 Probe insertion and placement in the drive body

This is the most delicate step of the procedure.

- Prepare the probe. Cut a piece of guide tubing (33 ga) and position it at the base of the probe (where the probe shank ends). This protective piece will be used to glue the probe to the spring, without hindering the mobility of the probe mechanism. The protective piece must not be glued to the shank at this stage.

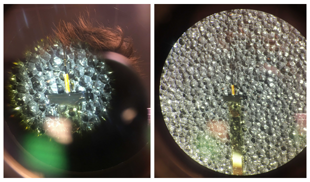

Figure 7: Close up of the protective guide tube positioned in the probe.

- Carefully hold the probe with forceps (from the black rectangle at the beginning of the shank) and insert it in the guide tube (the one left unglued) from above.
  **Note:** The distance from the protective tube on the probe and the guide tube on the body will determine the amount the probe can be moved. For stability reasons it is recommended to leave a maximum space of 2 mm.

- Carefully glue the protective tube on the probe to the bridge of the spring. To ensure a rapid procedure, apply accelerator to the bridge and glue to the protective tube on the probe. Then, gently move the probe in position and let the glue solidify. Once in position you should have the protective

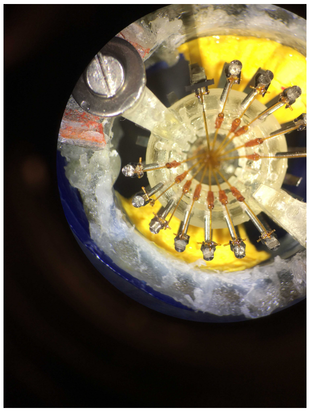

Figure 8: Tubing crown with silicon probe inserted and protective tube glued to the spring.

guide tubing glued to the bridge of the spring.
**Note:** At this stage you can still, carefully, move the probe up and down using forceps (from the black rectangle at the beginning of the shank).

### 1.9 Add the EIB

Once the silicon probe is firmly in place (1-2 hours of wait after gluing procedure), you can add the EIB.

- Add the EIB from above and screw it in place with 2 small screws. Alternatively you can use a piece of tubing to connect the EIB to the drive body.
- Before placing the EIB in its final position, make sure to leave approx. 2 mm between the EIB and the wings of the drive body. You can fill this space with a couple of pieces of tubing.
- Glue (with epoxy) the EIB to the wings of the drive body.
  **Note:** Make sure the EIB is firmly in place and it does not wobble. It could damage the probe otherwise.
- Once the EIB is firmly in place, connect the ZIF connector of the silicon probe to the ZIF port on the EIB.

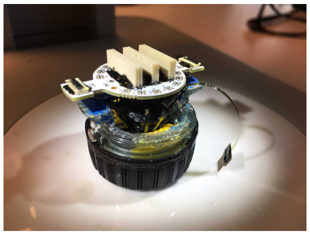

Figure 9: EIB glued to the drive body (probe not connected yet).

#### 1.10 Insert and wire tetrodes

Once the probe is well secured you can start a standard tetrode loading and wiring procedure (see Nguyen et al., 2009 for in-depth explanations). Insert tetrode into the shuttle tubes and cut the excess at the bottom of the guide array. Connect each electrode to the EIB using gold pins and glue the top of shuttle tubes with epoxy. In the proximity of the silicon probe pay more attention to the wiring procedure. In order to help reconstruct the final location of the tetrodes keep track of the mapping between the tetrodes position in the guide array and the EIB channels.

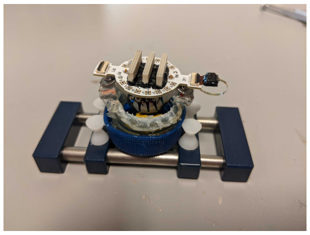

Figure 10: Drive in the holder for tetrode wiring, with the silicon probe connected via the ZIF connector.

#### 1.11 Tetrode cutting

Use serrated scissors.

- Before cutting tetrodes, move the silicon probe up. The lowest part of the shank (where elctrodes are) should be entirely back inside the tubing. This is to avoid damage to the shank of the silicon probe during the cutting procedure.
- Remove drive from holder.
- Place drive on the side with tetrodes parallel to the table with slight inclination to be able to visualise all tetrodes through the scope.

**Note:** In order to avoid any accidental damage to the flex-cable of the silicon probe, place the drive so that the flex cable is facing upwards.

- Hold scissors with serrated blade on top and smooth blade on the bottom.
- Cut tetrodes with a firm, single movement.
- The amount left will determine the total movement possible inside the brain. This varies depending on the target area. For the CA1 pyramidal layer 2.5 mm are enough.
- Add extra layer of epoxy between EIB and drive body to strengthen the structure.

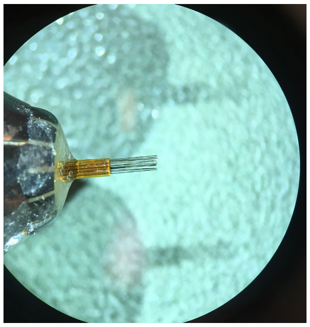

Figure 11: Tetrode ends at the bottom of the drive, cut at the desired length.

#### 1.12 Plating

- Plate the tetrodes.
  **Note:** At this stage the tip of the silicon probe is still inside the tubing structure.
- Once the tetrodes are plated you need to move them back inside the tubing structure of the array. You can carefully move them one by one using forceps (from the top, near the inner tube and EIB).
- Once all the tetrode tips are even with the bundle array you can glue the tetrodes in position. Add a drop of epoxy the the point at which the tetrodes exit the inner tube.

#### 1.13 Gluing the silicon probe in its final position

This is the final step before shielding. Once you glue the probe in position you will only be able to move it via the screw mechanism. After a few tests, we realized that it is safer (and more efficient) to set the silicon probe in its final position during drive building (rather than lowering it after the implant surgery).

- Gently lower the probe inside the tubing with forceps (push down from the black rectangle at the beginning of the shank).

- Measure, from the bottom of the drive, how much the silicon probe sticks out.

- Reach your desired length by moving up and down the probe.
  **Note:** In order to cover the entirety of CA1 you need to make the probe stick out of 2.2/2.3 mm.

- Glue with abundant epoxy the probe (at the level of the black rectangle at the beginning of the shank), to the protection guide tube.
  **Note:** Be careful not to spread glue along the silicon probe shank. It will hinder the mechanism.

### 1.14 Shielding

Shielding is the last step of the Hybrid Drive building procedure.

- You can use a cone shape plastic sheet to cover the body following the same shielding design of the flexDrive:
  `https://open-ephys.atlassian.net/wiki/spaces/OEW/pages/950325/6+-+Shielding+the+drive.`

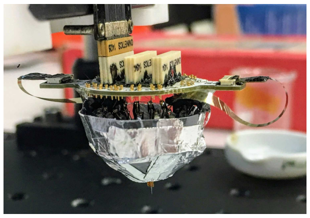

Figure 12: Hybrid drive with the body shield glued to the drive body.

- The lateral shields are important to protect the ZIF connectors.

- Mold the shape of the lateral protection using a small plastic cup.

- Cover this initial plastic protection with aluminium tape.

- Reinforce the structure by applying epoxy all over it.

- Position the cone over the ZIF connector and glue it to the body with epoxy.
  **Note:** It is important that probe ZIF connector does not touch the protection when you insert it. If your protection is too tight, repeated collision in the homecage (once the drive is implanted on the animal) will break this connection.

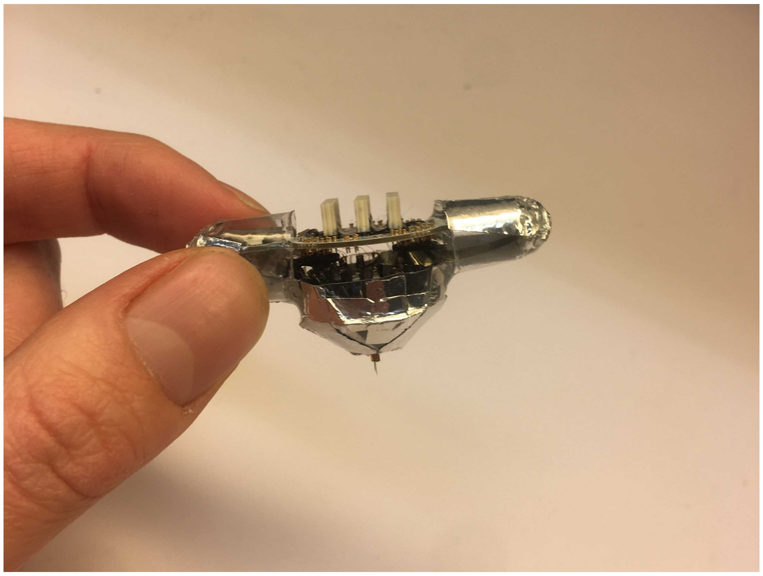

Figure 13: Hybrid drive with the lateral cone protections glued, ready for implant surgery.
